# Repurposing the aldose reductase inhibitor and diabetic neuropathy drug epalrestat for the congenital disorder of glycosylation PMM2-CDG

**DOI:** 10.1101/626697

**Authors:** Sangeetha Iyer, Feba S. Sam, Nina DiPrimio, Graeme Preston, Jan Verhejein, Kausalya Murthy, Zachary Parton, Hillary Tsang, Jessica Lao, Eva Morava, Ethan O. Perlstein

## Abstract

Phosphomannomutase 2 deficiency, or PMM2-CDG, is the most common congenital disorder of glycosylation affecting over 1,000 patients globally. There are no approved drugs that treat the symptoms or root cause of PMM2-CDG. In order to identify clinically actionable compounds that boost human PMM2 enzyme function, we performed a multi-species drug repurposing screen using a first-ever worm model of PMM2-CDG followed by PMM2 enzyme functional studies in PMM2-CDG patient fibroblasts. Drug repurposing candidates from this study, and drug repurposing candidates from a previously published study using yeast models of PMM2-CDG, were tested for their effect on human PMM2 enzyme activity in PMM2-CDG fibroblasts. Of the 20 repurposing candidates discovered in the worm-based phenotypic screen, 12 are plant-based polyphenols. Insights from structure-activity relationships revealed epalrestat, the only antidiabetic aldose reductase inhibitor approved for use in humans, as a first-in-class PMM2 enzyme activator. Epalrestat increased PMM2 enzymatic activity in four PMM2-CDG patient fibroblast lines with genotypes R141H/F119L, R141H/E139K, R141H/N216I and R141H/F183S. PMM2 enzyme activity gains range from 30% to 400% over baseline depending on genotype. Pharmacological inhibition of aldose reductase by epalrestat may shunt glucose from the polyol pathway to glucose-1,6-bisphosphate, which is an endogenous stabilizer and coactivator of PMM2 homodimerization. Epalrestat is a safe, oral and brain penetrant drug that was approved 27 years ago in Japan to treat diabetic neuropathy in geriatric populations. We demonstrate that epalrestat is the first small molecule activator of PMM2 enzyme activity with the potential to treat peripheral neuropathy and correct the underlying enzyme deficiency in a majority of pediatric and adult PMM2-CDG patients.

## Introduction

Deficiency of the enzyme phosphomannomutase-2 caused by loss-of-function mutations in the human *PMM2* gene was shown over two decades ago to be the basis of a recessive congenital disorder of glycosylation originally called CDG1 or CDG1a. The first clinical observation by Jaeken and colleagues of a “carbohydrate-deficient glycoprotein syndrome” occurred four decades ago (Jaeken et al, 1980). The researcher and patient communities now refer to the disease as PMM2-CDG, which is the most common congenital disorder of glycosylation affecting at least 1,000 patients worldwide (Chang et al, 2018). Classical pediatric clinical presentations include developmental delay, severe encephalopathy with axial hypotonia, abnormal eye movements, psychomotor retardation and cerebellar hypoplasia (Matthijs et al, 1997). As patients reach their teenage years and young adulthood, health challenges include hypogonadism, coagulation abnormalities and thrombotic events, retinitis pigmentosa and peripheral neuropathy (Monin et al, 2014) The prognosis for PMM2-CDG patients is poor and there is currently no FDA approved treatment that alleviates the symptoms of PMM2-CDG or any targeted therapy that safely increases PMM2 enzyme activity.

The PMM2 enzyme forms an obligate homodimer in the cytoplasm that converts mannose-6-phosphate to mannose-1-phosphate, an initial essential step in the *N*-linked glycosylation of proteins. Glucose-1,6-bisphosphate and mannose-1,6-bisphosphate are endogenous coactivators of PMM2 function, binding to and stabilizing PMM2 dimers (Andreotti et al, 2015). *N*-linked protein glycosylation is an evolutionarily conserved process that occurs in all animal cells throughout development and adulthood (Chang et al, 2018). PMM2-CDG is a multi-system, multi-organ disease because a minimal level of glycosylation is required at all times in all cells of the body, with cell types or organs more or less vulnerable to the complex sequelae of hypoglycosylation. Although a clear genotype-phenotype relationship is obscured by genetic and possibly environmental modifiers, as the residual level of PMM2 enzymatic activity increases the number and severity of organ systems affected decreases. For example, patients homozygous for a mutation in the promoter of PMM2 do not get PMM2-CDG or even a mild form of PMM2-CDG but instead have hyperinsulinemic hypoglycemia and polycystic kidney disease because this mutation impairs binding by a kidney- and pancreas-specific transcription factor to a chromatin loop in the promoter of PMM2 (Cabezas et al, 2017). As another example, hypoglycosylation of the calcium channel CACNA1A caused a gain-of-function channelopathy that in turn leads to an increase in stroke-like events in PMM2-CDG patients (Izquierdo-Serra et al, 2018).

Complete loss of *N*-linked protein glycosylation uniformly results in lethality of all animals in which PMM2 has been genetically knocked out, including humans. Homozygotes of the most common pathogenic variant, R141H, which is catalytically null, have never been observed alive in spite of the statistical predictions of population genetics (Matthijs et al, 1998; Kjaergaard et al, 1998). Those results indicate that there is a lower bound of PMM2 enzymatic activity (3-7%) required for viability. However the minimal PMM2 enzymatic activity above which disease is suppressed is unknown. Human genetics proves that this safety threshold varies from tissue to tissue and across stages of development. It further suggests that there are sharp tissue-specific transitions from physiology to pathophysiology with buffering capacity determined by both common and rare genetic modifiers in *N*-linked glycosylation and related metabolic pathways (Citro et al, 2019).

Over 80% of disease-causing PMM2 alleles are missense mutations resulting in amino-acid substitutions that destabilize an otherwise catalytically competent protein. Missense mutations fall into at least three biochemical classes: (i) protein destabilizing/misfolding mutations randomly distributed throughout the protein, (ii) dimerization defective mutations located in the monomer-monomer interface, and (iii) “catalytic dead” mutations in the active site (Yuste-Checa et al, 2015). Each PMM2 monomer forms a dimer with itself as a prerequisite for catalytic activity, though there need only be one functional active site per dimer (Andreotti et al, 2015). A plurality of PMM2-CDG patients across ethnic populations share the compound heterozygous genotype R141H/F119L, which pairs the aforementioned R141H null allele with the F119L dimerization-defective allele. Typically, patients present with R141H in compound heterozygosity with a hypomorphic missense allele, often a rare or private variants. Citro and colleagues argue that the unusual tolerance of PMM2 to missense mutations compared to every other CDG gene suggests a fitness advantage to being a PMM2 heterozygous carrier. As PMM2 enzymatic activity dips below 50%, at which point disease symptoms arise and how severely they arise are unknown because of the contribution of genetic modifiers that buffer the safety thresholds between health and disease. Similarly, it is unknown how small a boost in PMM2 enzymatic activity above disease baseline is required to elicit a therapeutic effect in PMM2-CDG patients.

A challenge for the PMM2-CDG community has been inviable mouse models and the lack of a gold standard cellular model of disease given the fact that *N*-linked protein glycosylation is a process that occurs in all cell types throughout life. A morpholino-knockdown model of PMM2-CDG in zebrafish displayed what appeared to be disease relevant phenotypes but these results have not been confirmed in a genetic knockout mutant (Cline et al, 2012). Fly models of PMM2-CDG were generated and also exhibited what appeared to be disease relevant phenotypes but they suffer from early larval lethality, which is a difficult phenotype to rescue in a genetic or chemical screen (Parkinson et al, 2016). For those reasons, Perlara PBC developed the first yeast models of PMM2-CDG (Lao et al, 2019).

Previously, we established evolutionarily conserved genotype-phenotype relationships across yeast and human patients between five PMM2 disease-causing mutations and their orthologous mutations in yeast. Overexpression of PMM2 in yeast cells lacking SEC53 (the yeast ortholog of PMM2) rescued lethality by mass action effects, and that there was no toxicity associated with overexpression of either PMM2 or SEC53 in yeast (Lao et al, 2019). Modest increases in SEC53 enzymatic activity translated into large phenotypic gains in yeast cell fitness. If the transition from physiology to pathophysiology is steep, is the transition from pathophysiology back to physiology equally steep? And are these sharp transitions conserved from yeast to humans? If the slopes of those transitions are conserved, small molecules are an attractive therapeutic modality not only because activation of PMM2 enzymatic activity appears to be well-tolerated but also because modest boosts in PMM2 enzymatic activity may be sufficient to produce real world clinical benefit and disease modification.

An academic drug discovery effort for PMM2-CDG involved expression of recombinant human PMM2 protein in bacteria, a primary target-based differential scanning fluorimetry screen, and a secondary fibroblast-based PMM2 enzymatic activity assay in order to identify pharmacological chaperones (Yuste-Checa et al, 2017). Although this approach proved the concept that it is possible to discover pharmacological chaperones and novel chemotypes that increase PMM2 enzymatic activity, only one early-stage tool compound was identified and it is far from clinical candidacy. Another approach proposed by Andreotti and colleagues is to increase the levels of glucose-1,6-bisphosphate (G16BP), an endogenous stabilizer and coactivator of PMM2 dimers (Monticelli et al, 2019). They propose that G16BP as a therapeutic pharmacological chaperone if it could be safely and effectively delivered to cells *in vivo*. We reasoned that rather than supplying exogenous chemically modified G16BP there may be small molecule drugs that pharmacologically increase endogenous G16BP by activating its synthesis or blocking its degradation or both. The biotech company Glycomine is developing an oral liposomal mannose-1-phosphate substrate replacement therapy that is currently in Phase 1.

In order to identify drug repurposing candidates that boost PMM2 enzyme function, we generated and characterized the first worm patient avatar of PMM2-CDG as an intermediate translational model situated between our previously published yeast models (Lao et al, 2019) and well-established PMM2-CDG patient fibroblasts. We created a viable hypomorphic loss-of-function F119L homozygous mutant worm for use in a primary *in vivo* phenotypic drug screen and a secondary *in vitro* worm PMM2 enzyme activity assay. We showed that this F119L hypomorphic mutant exhibits hypersensitivity to the proteasome inhibitor and ER stress inducer bortezomib, resulting in early larval arrest. Compounds that rescue arrested worm larvae and allow for normal development could activate ER stress response pathways to overcome chronic proteome-wide hypoglycosylation or boost PMM2 enzyme activity or boost PMM2 protein abundance. We adapted a low-throughput PMM2 enzymatic activity assay to a multi-well-plate assay using PMM2-CDG patient fibroblasts (Van Schaftingen & Jaeken, 1995) in order to determine which clinical candidates from model organism drug repurposing screens increase human PMM2 enzymatic activity.

The majority of repurposing candidates are “generally recognized as safe” plant-derived polyphenols, or phytochemicals, specifically a structurally-related group of antidiabetic and antioxidant dietary flavonoids. These flavonoids (e.gs., fisetin, ellagic acid) appear to share a common target and mechanism of action with a cinnamic acid derivative that specifically rescued a F119L yeast model of PMM2-CDG (Lao et al, 2019), namely aldose reductase inhibition. Based on structure-activity relationships, we found that epalrestat, a generic diabetic peripheral neuropathy drug and the only safe aldose reductase inhibitor approved for use in humans (Hotta et al, 2006), is a first-in-class PMM2 enzyme activator. The efficacy of epalrestat in four genotypically distinct PMM2-CDG fibroblasts tested in this study suggests that epalrestat could be given to PMM2-CDG patients who are compound heterozygous for R141H and any pathogenic variant. However, epalrestat treatment of PMM2-CDG fibroblasts did not increase PMM2 protein levels as measured by immunoblotting, suggesting that aldose reductase inhibition acts post-translationally to boost PMM2 enzyme activity. We propose that aldose reductase inhibition leads to an increase in glucose-1,6-bisphosphate levels as a result of glucose being shunted away from the polyol pathway toward production of phosphorylated glucose. Evidence from epalrestat-treated *pmm-2* mutant worms suggests there may also be a minor role for NRF2 as an indirect transcriptional activator of PMM2, which is consistent with a role for NRF2 activation in rescuing hypoglycosylation stress phenotypes a *pmm2* mutant zebrafish (Mukaigasa et al, 2018).

Epalrestat is an oral formulation of an aldose reductase inhibitor which has been commercially approved and available in Japan since 1992 for the treatment of diabetic neuropathies. It is also commercially available in India and China. The drug was never commercially approved in the United States or EU but it has a long and safe history. The drug’s ability to safely improve symptoms of neuropathy alone by reducing oxidative stress, increasing glutathione levels, and reducing intracellular sorbitol accumulation make it a desirable medication for PMM2-CDG patients who commonly suffer with various neuropathies. Based on the data presented herein, the newly discovered PMM2 enzyme activation mechanism of epalrestat may reduce the severity of the morbidities associated with PMM2-CDG.

## Materials and Methods

### Strains, cell lines and compounds

The *pmm-2* mutant VC3054 was obtained from the Caenorhabditis Genetics Center (CGC). It is a homozygous lethal deletion chromosome balanced by GFP-marked translocation. COP1626 is the *pmm-2* F125L (F119L in humans) hypomorphic homozygous mutant that was generated using CRISPR/Cas9 by NemaMetrix, Inc. The R141H/F119L patient fibroblast line GM20942 was obtained from Coriell. The R141H/E139K patient fibroblast line GM27386 was obtained from Coriell. The R141H/N216I patient fibroblast line was obtained from the Mayo Clinic biobank. The R141H/F183S patient fibroblast line was obtained from the Mayo Clinic biobank. Control fibroblasts GM05757, GM08398, GM08429 were ordered from Coriell. Screening was conducted using the 2,560-compound Microsource Spectrum library consisting of FDA approved drugs, bioactive tool compounds, and natural products. For all retests, compounds were reordered from Spectrum Discovery as 5 milligram dry powder stocks. Compounds were solubilized with fresh dimethylsulfoxide (DMSO) at high concentrations of 100mM or 50mM and stored as aliquots at -20C. For worm retesting, a 10mM stock was prepared.

### High-throughput larval growth assays in worms

The F125L/F125L homozygous *pmm-2* mutant growth screen was conducted in 384-well plates. 320 wells of the plate are filled with 25μM of compound (137.5nL) from the Microsource Spectrum library, dispensed into each well of the plate using an Echo 550 acoustic dispenser from Labcyte, Inc. In addition, all test wells were dispensed with 12.5nL of a 50mM stock of bortezomib solubilized in DMSO, resulting in a final concentration of 11μM. Control wells of the plate were filled with 150nL of DMSO (positive control), or 12.5nL of bortezomib and 137.5nL of DMSO (negative control). Following addition of test compounds, wells were dispensed with 5μL of bacterial media resuspended in S-medium containing cholesterol after adjusting the optical density to 0.45 at 600nm. Mutant animals were grown on standard nematode growth media agar plates until gravid. Adult worms were bleached using hypochlorite treatment to obtain eggs. Eggs were allowed to hatch overnight at 20°C in order to obtain synchronized L1 larvae. L1 larvae were filtered through 15-micron filters to ensure that the resulting population was clean before sorting. Fifteen L1 larvae are dispensed per well of the 384-well plate using a BioSorter large particle flow cytometer from Union Biometrica. Plates were sealed and incubated on a shaker at 20°C for five days. On the fifth day, plates were vortexed and spun down before the addition of 15μL of 8mM sodium azide. The addition of sodium azide immobilized worms after which they were imaged under transmitted light using a custom plate imager. Finally, automated image processing was run on each plate.

### Data analysis and statistical methods

We used the statistical program R to calculate Z-scores after processing the raw data, and we used Excel to calculate p values using t tests. We used custom image processing algorithms to extract areas occupied by worms per well. After outlier elimination among controls, i.e., elimination of data points on account of image or experimental artifacts, Z-scores were assigned to each well of the plate relative to the negative controls. Next, all wells that had a Z-score of greater than two in triplicate were isolated and manually inspected. Because bortezomib suppresses larval growth and induces arrest, we determined that a compound rescued the underlying defect if a well had a higher area occupied by worms relative to the negative control. Visual inspection often revealed that in primary screening positive wells, animals attained adulthood and produced progeny.

### PMM2 enzymatic assay in worms

L1-stage-synchronized F125L/F125L homozygous *pmm-2* mutant animals were grown (2,000 worms per plate) on NGM agar for 24 hours. After 24 hrs, worms were washed off plates into 15mL conical tubes, washed with filtered autoclaved water, pelleted at 3,200 rpm for 4 mins, and then resuspended in 2mL S-medium. HB101 bacteria grown in LB overnight was pelleted at 4,000 rpm for 10 minutes and resuspended in S-medium with cholesterol to an optical density of 0.35. For each experimental condition, 25mL of HB101 in S-medium was added to 50mL conical tubes along with 15,000 to 20,000 worms. Test compounds were dissolved in DMSO stock solutions and added to samples at a final concentration of 15µM. Samples were incubated at 20°C for 24 hours. After 24 hours, conical tubes were placed on ice, and worms were allowed to settle for 15 minutes. The bacterial supernatant was aspirated, and the worms were pelleted and washed with water. Worms were transferred to 1.5mL Eppendorf tubes, pelleted at 4°C, and lysed in 70-100µL homogenization buffer (20mM Hepes, 25mM KCl, 1mM DTT, 10µg/mL leupeptin, 10µg/mL antipain) on ice. The lysate was centrifuged, and 1-2µL of lysate was used for protein quantification using a Qubit. Lysate equivalent of 10µg protein was used per well to determine PMM-2 enzyme activity levels after adding the 200µL of assay buffer. Assay buffer without the substrate was used as the control and the assay was carried out for 3-4 hours with absorbance readings every 30 minutes at 340nm.

### PMM2 enzymatic assay development in R141H/F119L patient fibroblasts

Briefly, cells were seeded in a 96-well plate, homogenization buffer (20mM Hepes, 25mM KCl, 1mM DTT, 10µg/mL leupeptin, 10µg/mL antipain) was added, and plates were freeze-thawed at -80°C twice to lyse cells. Reaction buffer containing the substrate was then added to the wells of each plate. Plates were incubated at 37°C for 30 minutes and absorbance was read at 340nm at regular time points: 30, 60, 90, 120, 150, 180, 210, 240 and 270 minutes. All incubations are carried out with or without substrate (mannose-1-phosphate), and the difference between the two values was taken as the enzymatic activity. Enzyme activity was normalized to total lysate protein levels. Enzyme activity of wildtype fibroblasts and patient-derived compound heterozygous (F119L/R141H) fibroblast line were determined.

To assess if compounds from the phenotypic screens affected enzyme activity, all compounds were incubated with the mutant cell line at a concentration of 10μM for a 24-hour period. Following this, enzyme activity was assessed as described above. At least two biological replicates were conducted and enzyme activity in the presence of a test compound was compared to DMSO-treated mutant cell line. For ease of analysis, we compared the enzyme activity (as represented by NADPH concentration) of each treatment condition to activity of baseline, untreated mutant line at the last time point.

### Epalrestat treatments in optimized PMM2 enzymatic activity assay

Samples were lysed using sonication, after which total protein concentration was determined through Biorad LSR total protein detection assay. For each sample, 1.0 mg/mL protein wa used for further analysis. After lysis, samples were centrifuged at 10,000 rpm for 10 minutes. After centrifugation, 80 µL of supernatant was transferred to a microtiter plate. PMM2 substrate mannose 1-phosphate (Man1P) was added to each sample after which the plate was transferred to a FLUOstar Omega Plate Reader to determine fluorescent excitation after 30 and 40 minutes as a measure of PMM2 enzyme activity. All samples were run in duplicate with each run including healthy control and disease control standards.

Enzyme activity scores are presented as nmol/h/mg protein using the following formula:

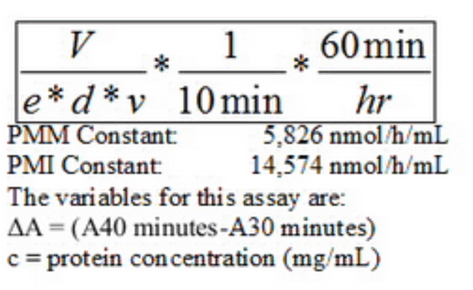

### Keap1-NRF2 activation luciferase reporter assay in U2OS cells

Hit compounds were tested by DiscoverX (San Diego) using the PathHunter® eXpress Keap1-NRF2 Nuclear Translocation Assay. Briefly, PathHunter cells are plated and incubated for 24 hours at 37°C. 10µL of test compound was added and cells were incubated with compound for 6 hours at room temperature. Working detection reagent solution was added and plates were incubated for 60 minutes at room temperature. Chemiluminescence signal was read by a SpectraMax M3. Methyl CDDO ester was used as the positive control. To be called a NRF2 activator, a hit compound had a half-maximal effective concentration (EC_50_) less than 10µM and a slope comparable to methyl CDDO ester.

### PMM2 protein quantification by immunoblotting in fibroblasts

To isolate protein, cells were washed in 1X PBS and pelleted. Cell pellets were solubilized in RIPA Buffer + Protein Inhibitor Cocktail. Protein concentration was determined using the BCA method. Immunoblotting was performed as follows. 30µg cell protein was separated on a 10% Bis-Tris gel (NP0301PK2, ThermoFisher, Inc) and blotted onto nitrocellulose membranes using the standard protocol provided by the manufacturer. Rabbit polyclonal antibody against human PMM2 (10666-1-AP, Proteintech) was diluted 1/333 (30µL in 10mL blocking buffer). Mouse monoclonal antibody against human ACTB (AC004, CiteAb) was diluted 1:15,000 (0.66µL in 10mL blocking buffer). Membranes were blocked for 30 minutes at 4°C in SEA Block blocking buffer (37527, ThermoFisher. Primary antibodies were diluted in 10mL SEA Block. Membranes were incubated on a rocker in primary antibody dilution overnight at 4°C. Membranes were washed six times for 10 minutes in 40mL 1X PBS+ 0.1% Tween at room temperature. Donkey anti-mouse green (SA5-10172, ThermoFisher) and donkey anti-rabbit red (SA5-10042, ThermoFisher) fluorescent antibodies were diluted 1:5,000 (2µL each in 10mL blocking buffer). Membranes were incubated in secondary antibody dilution for one hour at 4°C. Membranes were then washed six times for 10 minutes in 40mL 1X PBS+ 0.1% Tween at room temperature. Finally, membranes were washed once for 10 minutes in 40mL 1X PBS at room temperature and immediately visualized.

### Quantitative RT-PCR of worms

Worms were age-synchronized and collected as day 1 adults. Worm pellets were homogenized and RNA extracted following published protocols (Guthmueller et al, 2011). ER stress markers were selected based on the *pmm2* zebrafish model (Mukaigasa et al, 2018). Cq stands for cycle quantification.

## Results

### Generation and phenotyping of the first nematode model of PMM2-CDG

We addressed the lack of animal models for PMM2-CDG that are amenable to unbiased high-throughput phenotypic screens by creating a series of worm patient avatars. The F52B11.6 gene sequence in *C. elegans* is orthologous to human PMM2. Nematode *PMM-2* shares 54% identity with the human protein, and the two most common disease-causing mutations -- R141H and F119L -- are evolutionarily conserved. We first characterized a heterozygous *pmm-2* null strain. This strain has a 518-base-pair deletion in the *PMM-2* open reading frame. This deletion strain is documented to be larval lethal, which we confirmed (data not shown). The heterozygous animals produce few homozygous null progeny, insufficient for the scale of a high-throughput drug screen. We used quantitative RT-PCR to measure the amount of worm *PMM-2* mRNA expression in *pmm-2* heterozygote null animals. We observed that worm *PMM-2* mRNA levels are the same in wildtype N2 animals and *pmm-2* heterozygote null animals, indicating that this mutant compensates for hemizygosity at the transcriptional level by boosting *PMM-2* expression to levels normally expressed in wildtype worms with two functional copies of *PMM-2* (**Supplementary Figure 1**). Because *pmm-2* homozygous nulls are inviable and *pmm-2* heterozygous nulls do not exhibit overt growth and developmental delay phenotypes, we engineered a novel *pmm-2* hypomorphic mutant strain using CRISPR/Cas9 and examined it for phenotypes amenable to a high-throughout image-based growth assay and a low-throughput whole-animal PMM2 enzyme activity assay.

We created a homozygous F125L/F125L missense mutant, as worm F125 is orthologous to human F119. F119 sits in the dimer interface and the F119L mutation results in a defect in dimerization. The homozygous *pmm-2* F125L (hereafter referred to as F119L) strain is homozygous viable and does not exhibit larval lethality, growth defects or any observable locomotor defects in liquid media. We verified that worm *PMM-2* mRNA transcript levels in the F119L mutant are comparable to wildtype N2 animals (**Supplementary Figure 1**). In order to determine if the F119L homozygote mutant is a bona fide model of PMM2-CDG, we measured worm PMM2 enzyme activity in F119L homozygotes after whole-animal lysis. As expected, this mutant has reduced PMM2 enzymatic activity, while by contrast PMM2 enzymatic activity is unchanged in a *png-1* homozygous null mutant and model of NGLY1 Deficiency, another glycosylation disorder, and wildtype N2 worms (**Figure 1**).

**Figure 1.**
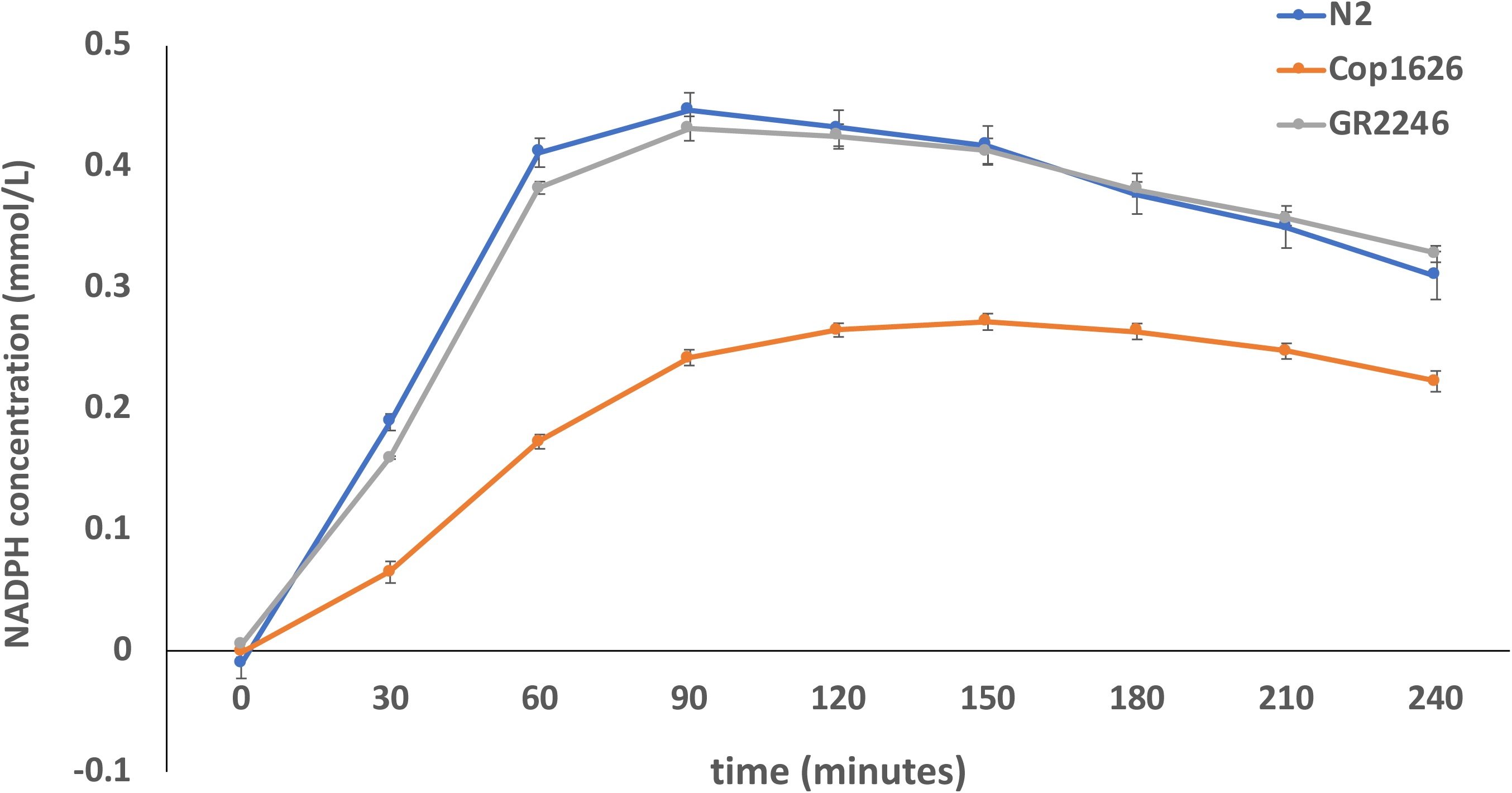
Plot of PMM2 enzymatic activity in whole-animal lysates of *pmm-2*^F126L^ mutant worms. Phosphomannomutase activity is measured by production of NADPH concentration (mmol/L) over time. The blue line indicates phosphomannomutase activity in wildtype N2 worms. The orange line indicates phosphomannomutase activity in COP1626 worms (*pmm-2*^F125L^ homozygote mutant). The gray line indicates phosphomannomutase activity of GR2246 worms, a *png-1* deletion mutant which serves as a negative control. The same amount of protein was used in lysates from each strain.

### Drug repurposing screen using *pmm-2* F125L/F125L mutant worms

Because the F119L homozygous mutant strain presented no constitutive growth and developmental timing differences compared to wildtype worms, it was not amenable to high-throughput screening as is. We tested if *pmm-2* mutant animals are more sensitive to pharmacological stressors that disrupt proteasomal processes and induce ER stress. Literature evidence shows that tunicamycin (Buzzi et al, 2011) and bortezomib (Tillman et al, 2018) exposure causes proteasomal stress, activates the unfolded protein response and induces larval arrest in worms. We treated F119L mutant animals with increasing concentrations of tunicamycin and bortezomib. We sought to establish if either compound affected growth and development of mutant animals at concentrations lower than similarly exposed wildtype worms. Tunicamycin did not produce differential larval arrest in mutants and wildtype worms possibly because the F119L mutation is not sensitizing enough (data not shown). It is possible tunicamycin might produce a differential larval arrest in the context of modeling a more severe *PMM2* variant in worms.

As shown in **Figure 2**, we found that *pmm-2* mutant worms exposed to bortezomib, a reversible proteasome inhibitor, displayed larval arrest at concentrations more than 10-fold lower than wildtype. At the highest concentration tested (13.6µM), F119L mutant worms uniformly arrested as small larvae with morphological defects or were dead, whereas wildtype worms at this dose were viable late-stage larvae or young adults (**Figure 1A**). Starting at 1.6µM bortezomib, the size distribution of *pmm-2* mutant worms separates from the size distribution of wildtype worms. At concentrations higher than 6.8µM, *pmm-2* mutant worms arrest as small sick larvae with no overlap between mutant and wildtype worm size distributions (**Figure 2B**). We decided to screen the Microsource Spectrum drug repurposing library in triplicate for compounds that rescue early larval arrest in the presence of 11μM bortezomib, a concentration which guaranteed robust statistical separation of positive control Z-scores (DMSO-treated *pmm-2* mutant worms) from negative control Z-scores (bortezomib-treated *pmm-2* mutant worms). We previously screened the Microsource Spectrum library using yeast models of PMM2-CDG (Lao et al, 2019). We anticipated identifying overlapping chemotypes active in both species.

**Figure 2.**
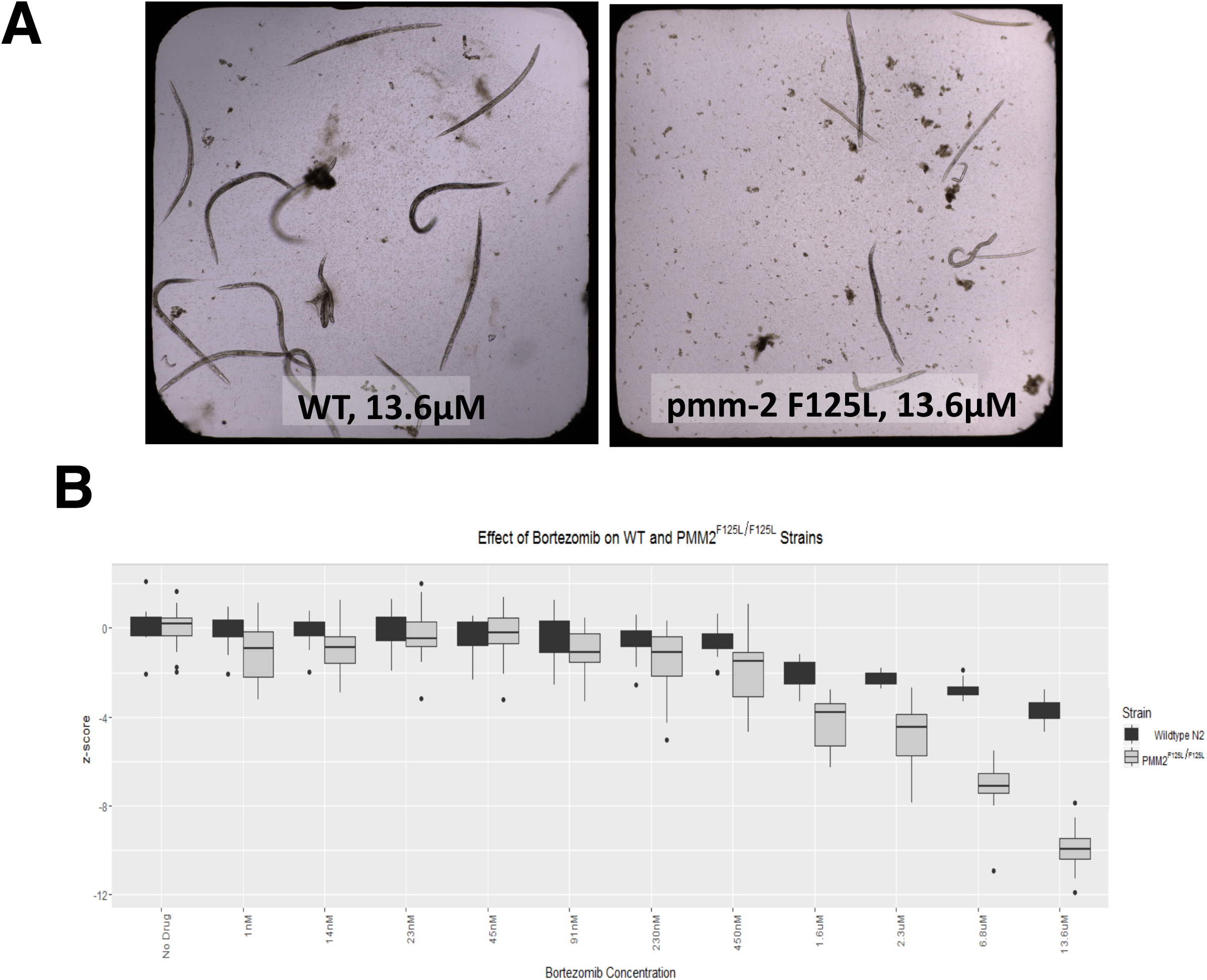
*pmm-2* F125L homozygote mutant worms are more than 10 times more sensitize to bortezomib than wildtype worms. (**A**) Side-by-side comparison of a representative well of a clear-bottom 384-well plate containing wildtype worms (left) and a representative well of a clear-bottom 384-well plate containing *pmm-2* F125L/F125L worms treated with 13.6µM bortezomib, the highest dose tested. (**B**) Z-score box and whisker plot comparing *pmm-2* F125L/F125L worms (gray boxes) versus wildtype worms (black boxes) treated with the following range of concentrations of bortezomib: 1nM, 14nM, 23nM, 45nM, 91nM, 230nM, 450nM, 1.6µM, 2.3µM, 6.8µM, 13.6µM. Z-score labels on the y-axis are 0, -4, -8 and -12. Individual black dots represent statistical outliers.

Representative positive control wells and negative control wells are shown in **Figure 3**. Note that the worms in negative control wells are still viable even if they are arrested (**Figure 3B**). Statistically significant separation between positive and negative controls allowed us to identify both suppressors and enhancers, which include toxic compounds (**Supplementary Figure 2**). We only examined suppressors in the context of this study, i.e., compounds that fully rescue early larval arrest induced by 11µM bortezomib. Note that the worms in wells containing suppressors look indistinguishable from worms in positive control wells (**Fig. 3D** versus **Fig. 3B**). 20 compounds had Z-scores greater than five in all three replicates (**Figure 3A**), resulting in a suppressor hit rate of 0.781%, which is consistent with the hit rates we observed in previous whole-organism phenotypic drug repurposing screens in flies (Rodriguez et al, 2018) and yeast (Lao et al, 2019).

**Figure 3.**
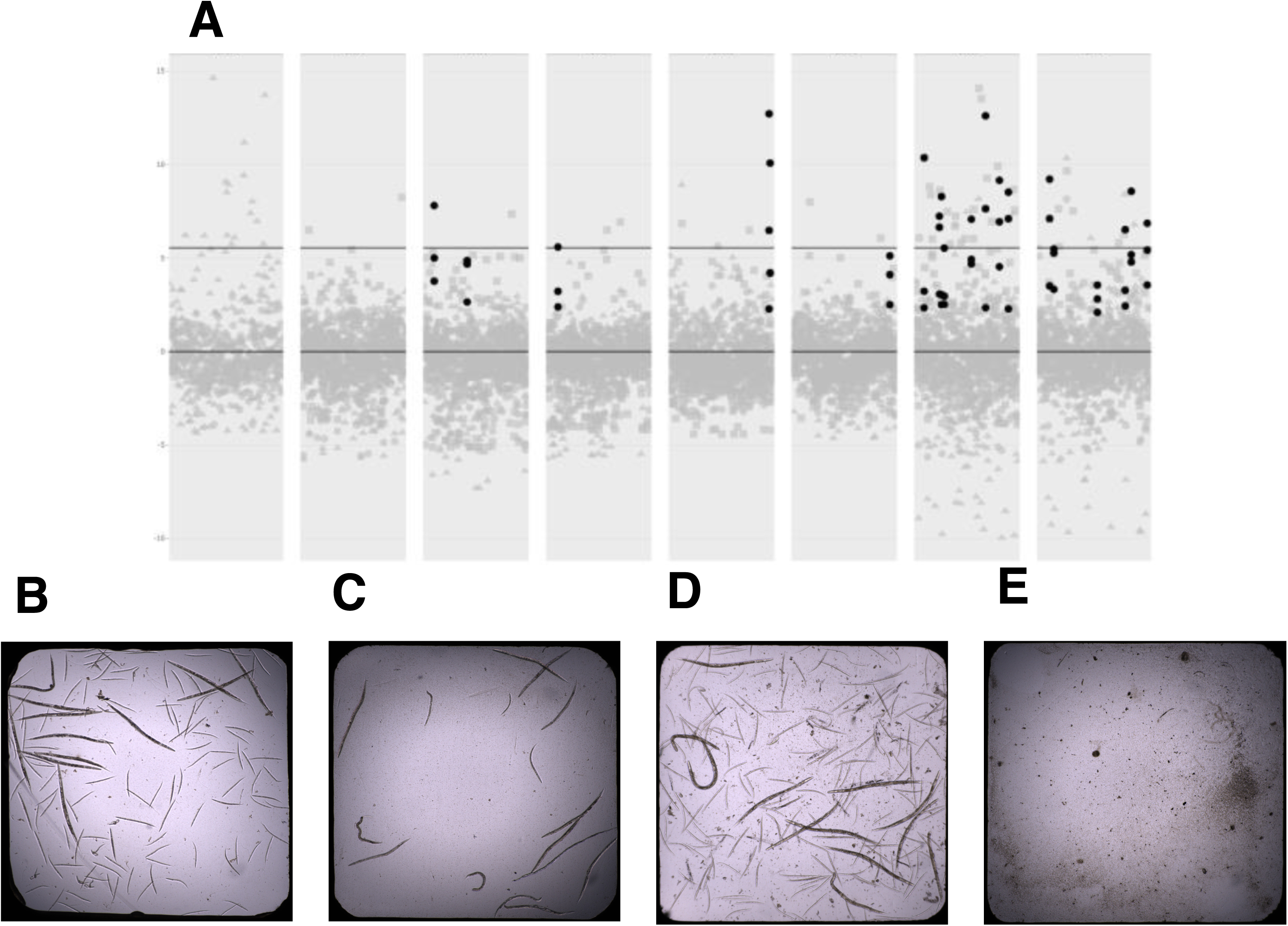
Summary of drug repurposing screen of *pmm-2* F125L/F125L mutant worms. (**A**) Three replicates of the Microsource Spectrum library screen (replicate 1 = gray circles, replicate 2 = gray squares, replicate 3 = gray triangles). Black circles represent the 20 hit compounds with Z-scores greater than five in all three replicates. Each column represents one of the eight library plates. The lower black line indicates Z-score = 0. The upper black line indicates Z-score = 5. (**B**) Image of a representative positive control well into which L1 DMSO-treated *pmm-2* F125L/F125L mutant worms were dispensed and incubated at 20°C for 5 days. 12 adults are visible along with their progeny. (**C**) Representative negative control well into which L1 11µM-bortezomib-treated *pmm-2* F125L/F125L mutant worms were dispensed and incubated at 20°C for 5 days. 16 animals are visible ranging from L1-L4 larval stages. (**D**) Representative suppressor (hit) well into which L1 *pmm-2* F125L/F125L mutant worms were dispensed and incubated at 20°C for 5 days. 12 adults are visible along with their progeny. (**E**) Representative enhancer/toxic well into which L1 11µM-bortezomib-treated *pmm-2* F125L/F125L mutant worms were dispensed and incubated at 20°C for 5 days. No living larvae are visible.

### Repurposing candidates are antidiabetic and antioxidant phytochemicals

12/20 (60%) of the repurposing candidates (suppressors) are generally recognized as safe (GRAS) dietary polyphenols found in fruits, vegetables, roots, spices and trees – specifically aglycone flavonoids and flavonoid glycosides: fisetin, rhamnetin, pyrogallin, purpurogallin-4-carboxylic acid, quercetin tetramethyl ether, ellagic acid, gossypetin, hieracin (tricetin), baicalein, koparin, epicatechin monogallate and theaflavin monogallate. 3/20 (15%) of the hit compounds are the simple building blocks or scaffolds of more complex polyphenols, and for that reason were not further considered: 3-methoxycatechol; 2,3,4-trihydroxy-4-methoxybenzophenone; 3,4-didesmethyl-5-deshydroxy-3-ethoxyschleroin. 3/20 (15%) are catecholamines: levodopa, ethylnorepinephrine, and dobutamine. Levodopa is a precursor to dopamine. Ethylnorepinephrine is a sympathomimetic drug and bronchiodilator. Dobutamine is a β1-adrenergic receptor agonist approved for the treatment of heart failure. The remaining two chemically simple compounds may be classified as nonspecific antioxidants: amidol and edavarone.

Plant-based polyphenols have complex polypharmacology, meaning they are biologically active at multiple protein and nonprotein targets in the cell. As a result, plant-based polyphenols have multiple mechanisms of action depending on the concentration tested and length of treatment in a diverse array of *in vitro* and *in vivo* models. What unites many dietary phytochemicals are their antioxidant and anti-inflammatory properties, which in human physiology translates to known antidiabetic (hypoglycemic) and potential senolytic effects.

Pyrogallin contains a tropolone moiety and is the decarboxylated derivative of purpurogallin-4-carboxylic acid. Fisetin, gossypetin, hieracin, baicalein and koparin are all structurally related flavones differing in number and location of hydroxyl groups. Quercertin tetramethyl ether and rhamnetin are O-methylated derivatives of quercetin, the most consumed dietary flavonoid. Ellagic acid results from the hydrolysis of more complicated tannins. Epicatechin monogallate and theaflavin monogallate, which contains the tropolone core scaffold that is pyrogallin, are flavanols present in green and black teas. It is known that gossypetin can directly inactivate bortezomib, resulting in false positive hits so we removed it from further consideration (Glynn et al, 2015).

The chemical structures of all twenty PMM2 repurposing candidates are shown in **Figure 4**. Remarkably, our flavonoid repurposing candidates have all been reported to be aldose reductase inhibitors (ARI) in the single to low double digit µM concentration range, and therefore would display this activity under the conditions of the primary drug repurposing screen and secondary hit validation assays. Fisetin, quercetin tetramethylether and rhamnetin were shown to inhibit aldose reductase with half-maximal inhibitory concentrations of 3.7µM, 25µM and 2.7µM, respectively (Matsuda et al, 2002). Quercetin was identified in 1975 as an aldose reductase inhibitor (Varma et al, 1975; Varma et al, 1977).

**Figure 4.**
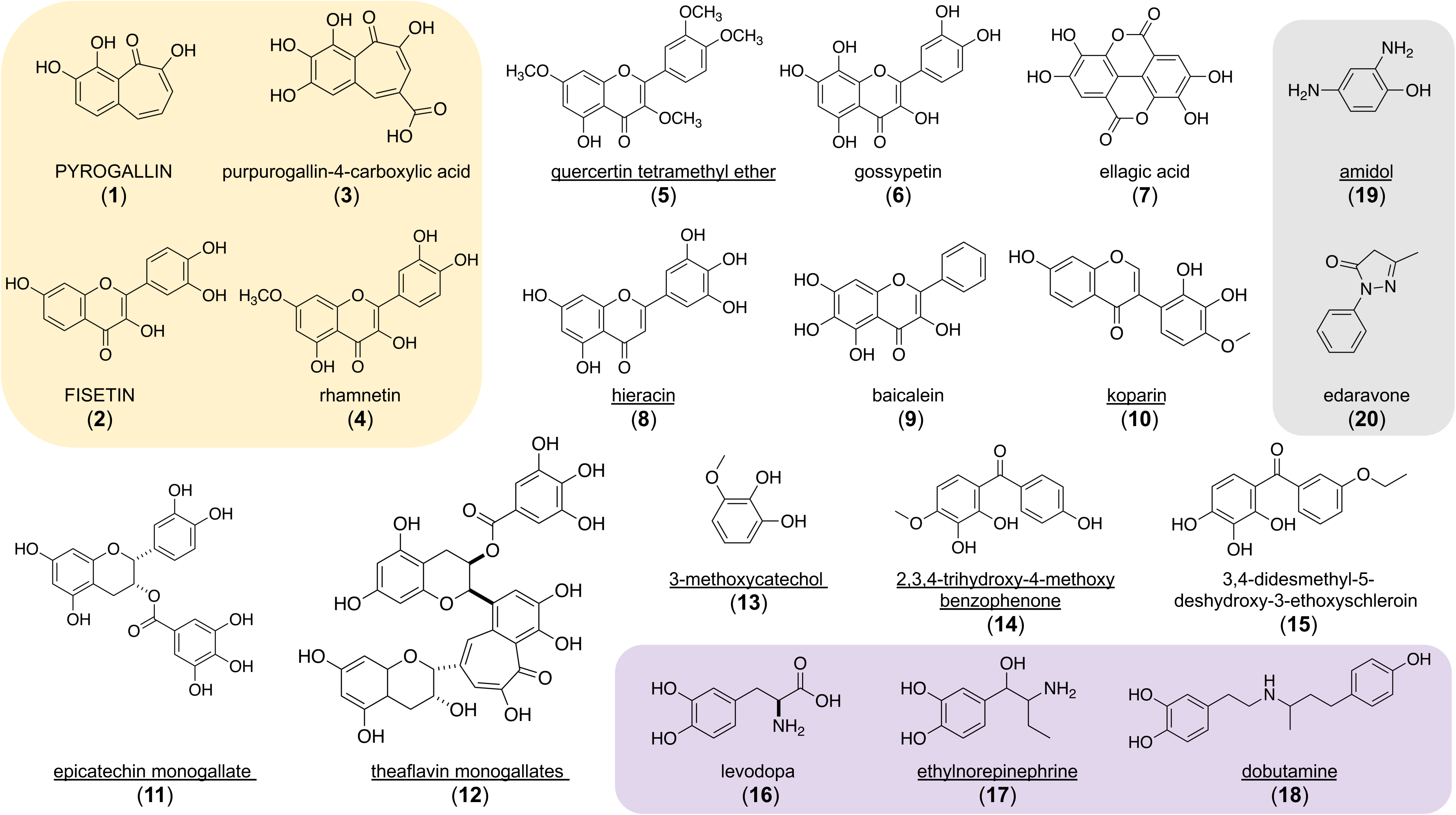
Chemical structures of 20 drug repurposing candidates discovered in a worm *pmm-2* bortezomib chemical modifier screen. Fully capitalized compounds are active in the human PMM2 enzyme activity assay performed in R141H/F119L PMM2-CDG fibroblasts as described in *Materials and Methods*. Underlined compounds are active in at least one of three yeast PMM2-CDG models described in Lao et al, 2019. The yellow box indicates compounds that are active in the Keap1-Nrf2 cellular reporter assay described in *Materials and Methods*. The purple box indicates compounds that are catecholamines. The grey box indicates structural singletons with non-selective antioxidant properties. (**1**) pyrogallin. (**2**) fisetin. (**3**) purpurogallin-4-carboxylic acid. (**4**) rhamnetin. (**5**) quercetin tetramethyl ether. (**6**) gossypetin. (**7**) ellagic acid. (**8**) hieracin (tricetin). (**9**) baicalein. (**10**) koparin. (**11**) epicatechin monogallate. (**12**) theaflavin monogallate. (**13**) 3-methoxycatechol. (**14**) 2,3,4-trihydroxy-4-methoxybenzophenone. (**15**) 3,4-didesmethyl-5-deshydroxy-3-ethoxyschleroin. (**16**) levodopa. (**17**) ethylnorepinephrine. (**18**) dobutamine. (**19**) amidol. (**20**) edaravone.

The flavonoid 2’-2’-bisepigallocatechin digallate (**Supplementary Figure 3**) was identified as one of three repurposing candidates in our previously published yeast PMM2 drug repurposing study (Lao et al, 2019), and is structurally related to theaflavin digallate. This result demonstrates that specific plant-based polyphenols rescue in both yeast and worm species paradigms by a conserved mechanism of action. Plant-based polyphenols also are all well-known antioxidants that activate cytoprotective responses like the Keap1-Nrf2 pathway, which is not conserved in yeast. We tested the 20 worm repurposing candidates in yeast PMM2-CDG models and in a human cell-based Keap1-Nrf2 activity assay.

### Cross-species retesting of worm repurposing candidates in yeast and fibroblasts

We performed two secondary screens in order to prioritize the 20 worm repurposing candidates for final validation testing in PMM2-CDG patient fibroblasts in a human PMM2 enzyme activity assay representing the most common genotype R141H/F119L. First, we tested all 20 worm repurposing candidates in three genotypically distinct yeast PMM2-CDG models (Lao et al, 2019). 10/20 (50%) of the worm repurposing candidates improved growth of one or more yeast PMM2 models (**Figure 4; Supplementary Figure 7**). Theaflavin monogallate, epicatechin monogallate and koparin and ethylnorepinephrine rescued two out of three yeast PMM2-CDG models but were not considered further. Quercetin tetramethylether, amidol and dobutamine rescue growth in a dose-dependent manner in all three yeast PMM2-CDG models. Quercetin tetramethylether had the largest effect size, so we began to suspect that it acts as an aldose reductase inhibitor because aldose reductase is conserved in yeast.

It is possible that inactivity in the yeast PMM2-CDG models could be the result of compound insolubility or instability in yeast media versus in worm media, or other organism-specific assay variables. In order to dissect the polypharmacology of plant-based polyphenols, we tested all 20 worm repurposing candidates in a Keap1-Nrf2 reporter human cell assay. We reasoned that plant-based polyphenols are known antioxidants, and some have been reported to activate the transcriptional regulator NRF2. Compounds that scored positively in the Keap1-Nrf2 reporter assay were then tested in a PMM2 enzyme activity assay on PMM2-CDG R141H/F119L fibroblasts in order to triage the list of compounds down to one or a few most promising repurposing candidates. We optimized PMM2 enzymatic assays for worms and for patient fibroblasts based on the previously reported protocol (Van Schaftingen & Jaeken, 1995).

When we tested all 20 worm repurposing hits in a Keap1-Nrf2 reporter assay in human cells, only 4/20 (20%) of the compounds activated NRF2: fisetin, rhamnetin, pyrogallin and purpurogallin-4-carboxylic acid (**Figure 4**). Fisetin is a known NRF2 activator (Smirnova et al, 2011). The other three were not previously known to activate NRF2. None of the four worm repurposing candidates that activate the Keap-Nrf2 reporter assay rescued any of the yeast PMM2-CDG models, which is the expected result given the lack of conservation of NRF2 in yeast. Pyrogallin, purpurogallin-4-carboxylic acid, rhamnetin and fisetin do not rescue any of the yeast PMM2-CDG models tested. In fact, all four inhibit growth of all three yeast PMM2-CDG yeast models tested, with pyrogallin and purpurogallin-4-carboxylic acid having larger effect sizes than fisetin and rhamnetin (**Supplementary Figure 7**).

### Validating aldose reductase inhibitors in PMM2 enzymatic assays in worms and fibroblasts

Next, we tested the four worm repurposing candidates that activated the Keap1-Nrf2 pathway in a human PMM2 enzymatic activity assay using a R141H/F119L PMM2-CDG patient fibroblast line. We also tested the three yeast PMM2 repurposing candidates reported in Lao *et al* whose structures are shown in **Supplementary Figure 3**. Of those seven compounds, only the yeast repurposing candidate alpha-cyano-4-hydroxycinnamic acid, or CHCA, robustly activated PMM2 enzymatic activity not only in PMM2-CDG R141H/F119L fibroblasts but also in the F119L mutant worm (**Figure 5**; chemical structure shown in **Supplementary Figure 3**). 15µM CHCA increased worm PMM2 enzymatic activity by 50% over baseline but the variance is large so the result is not statistically significant (p = 0.4175). Similarly, 10µM CHCA increased human PMM2 enzymatic activity by 40% and is marginally statistically significant because of the variance (p = 0.0436) as shown in **Figure 5B**. The worm repurposing candidates pyrogallin, purpurogallin-4-carboxylic acid and fisetin activated human PMM2 enzymatic in PMM2-CDG R141H/F119L fibroblasts (**Figure 4**) but they inhibit all three yeast PMM2-CDG models at all three concentrations tested. Therefore, we deprioritized those three compounds and focused on CHCA because it had the largest effect size in terms of increasing PMM2 enzymatic activity.

**Figure 5.**
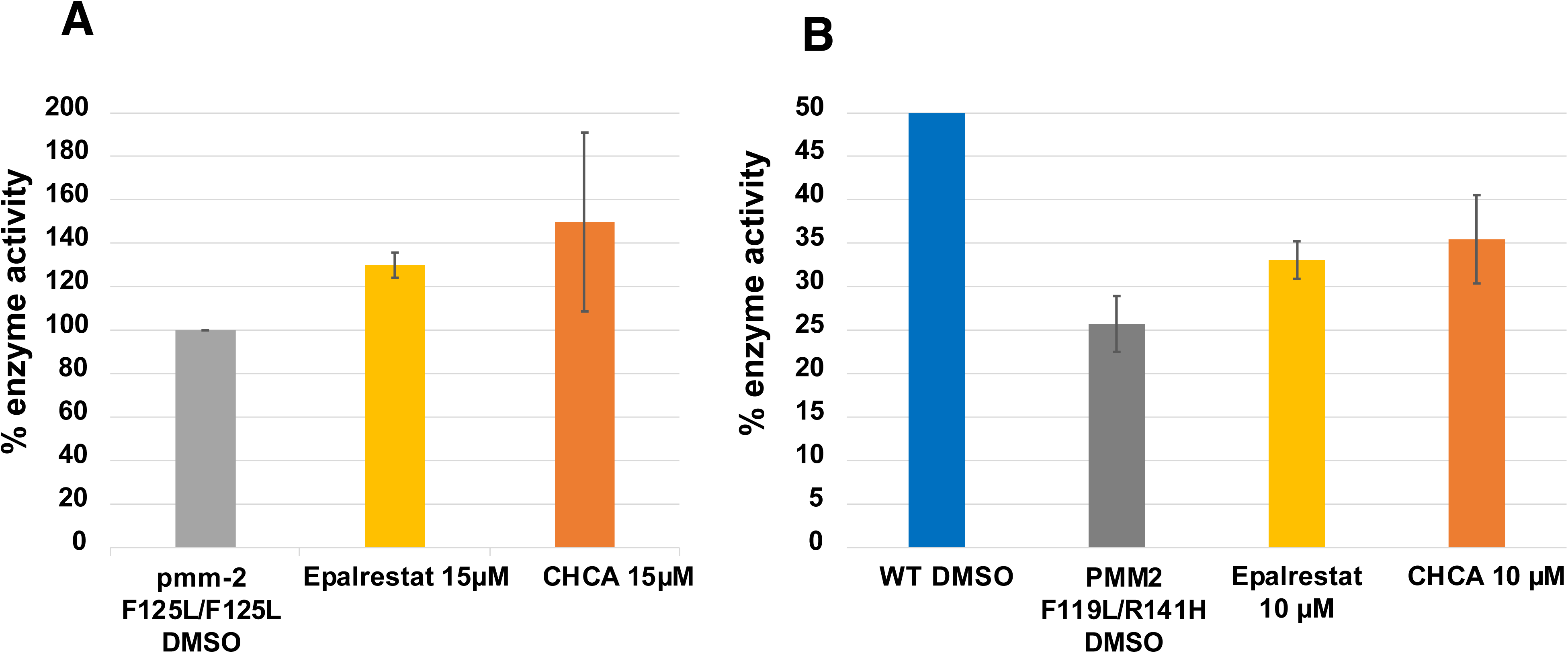
Aldose reductase inhibitors alpha-cyano-4-hydroxycinnamic acid (CHCA) and epalrestat increase PMM2 enzymatic activity in worms and patient fibroblasts. (**A**) Y-axis is percentage of PMM2 enzymatic activity normalized to the *pmm-2* F125L/F125L (F119L) homozygote mutant treated with DMSO vehicle. From left to right:; *pmm-2* F125L/F125L (F119L) homozygote mutant treated with 15µM epalrestat for 24 hours (p = 0.0352); *pmm-2* F125L/F125L (F119L) homozygote mutant treated with 15µM CHCA for 24 hours (p = 0.4175). (**B**) Y-axis is percentage of PMM2 enzymatic activity relative to wildtype (100%). 50% represents the activity level of unaffected heterozygous carriers. From left to right: GM20942 R141H/F119L PMM2-CDG patient fibroblasts treated with 10µM CHCA (p = 0.0436); GM20942 R141H/F119L PMM2-CDG patient fibroblasts treated with 10µM epalrestat (p = 0.00825); GM20942 R141H/F119L PMM2-CDG patient fibroblasts treated with DMSO vehicle; wildtype control fibroblasts treated with DMSO vehicle. The same amount of protein was used in lysates from each strain. Error bars represent standard error. P values were determined by unpaired t test (T.TEST function in Excel).

CHCA is a potent aldose reductase inhibitor (Zhang et al, 2016), which is consistent with the known ARI mechanism of action of dietary flavonoids. The predominant ARI pharmacophore contains a terminal carboxylic acid moiety like CHCA. We reasoned that the variance associated with CHCA may be due to its relative pharmacological promiscuity and off-target effects. Therefore, we tested the following nine commercially available aldose reductase inhibitors (ARIs) representing the known ARI pharmacophores, including those containing the terminal carboxylic acid moiety shared with CHCA: tolrestat, ranirestat, imirestat, zopolrestat, sorbinil, ponalrestat, alrestatin, fiderastat and epalrestat. Tolrestat, imirestat, ponalrestat and alrestatin increased PMM2 enzymatic activity in only one of three replicates (data not shown).

Of the ARIs tested, only carboxylic acid-containing epalrestat and CHCA reproducibly increased PMM2 enzymatic activity in both worms (**Figure 5A**) and fibroblasts (**Figure 5B**). In *pmm-2* F119L mutant worms, 15µM epalrestat treatment caused a 30% increase in PMM2 enzymatic activity (p = 0.0352). In R141H/F119L PMM2-CDG fibroblasts, 10µM epalrestat also caused a 30% increase in PMM2 enzymatic activity over baseline (p = 0.00825). Epalrestat happens to be the only ARI that is orally bioavailable, brain-penetrant, well-tolerated and approved for use in humans. Epalrestat was approved for the treatment of diabetic neuropathy in geriatric patients in Japan in 1992 (Hotta et al, 1996), and is also available in China and India, but has not been approved by the Food and Drug Adminstration (FDA) for any indications. In light of all these data, we deprioritized CHCA, which is not approved for use in humans, in favor of the safe and generic epalrestat that has been used for decades in Asia.

### Epalrestat boosts PMM2 enzyme activity in multiple PMM2-CDG patient fibroblasts

We tested whether the ARI epalrestat increases human PMM2 enzymatic activity in more than one PMM2 genotype and genetic background. Even though the compound heterozygous genotype R141H/F119L is the most common genotype observed in PMM2-CDG patients worldwide, there is a long tail of private missense mutations *in trans* with R141H so we tested three additional genotypes: R141H/E139K, R141H/N216I and R141H/F183S. Incubation of human PMM2-CDG derived fibroblasts with 10µM epalrestat for 24 hours led to an increase in PMM2 enzymatic activity in all tested samples **(Figure 6)**. As summarized in **Table 1**, each PMM2-CDG fibroblast line has a different PMM2 enzymatic activity baseline, reflecting the severity of the respective PMM2 mutations correlating with enzyme activity impairment.

**Figure 6.**
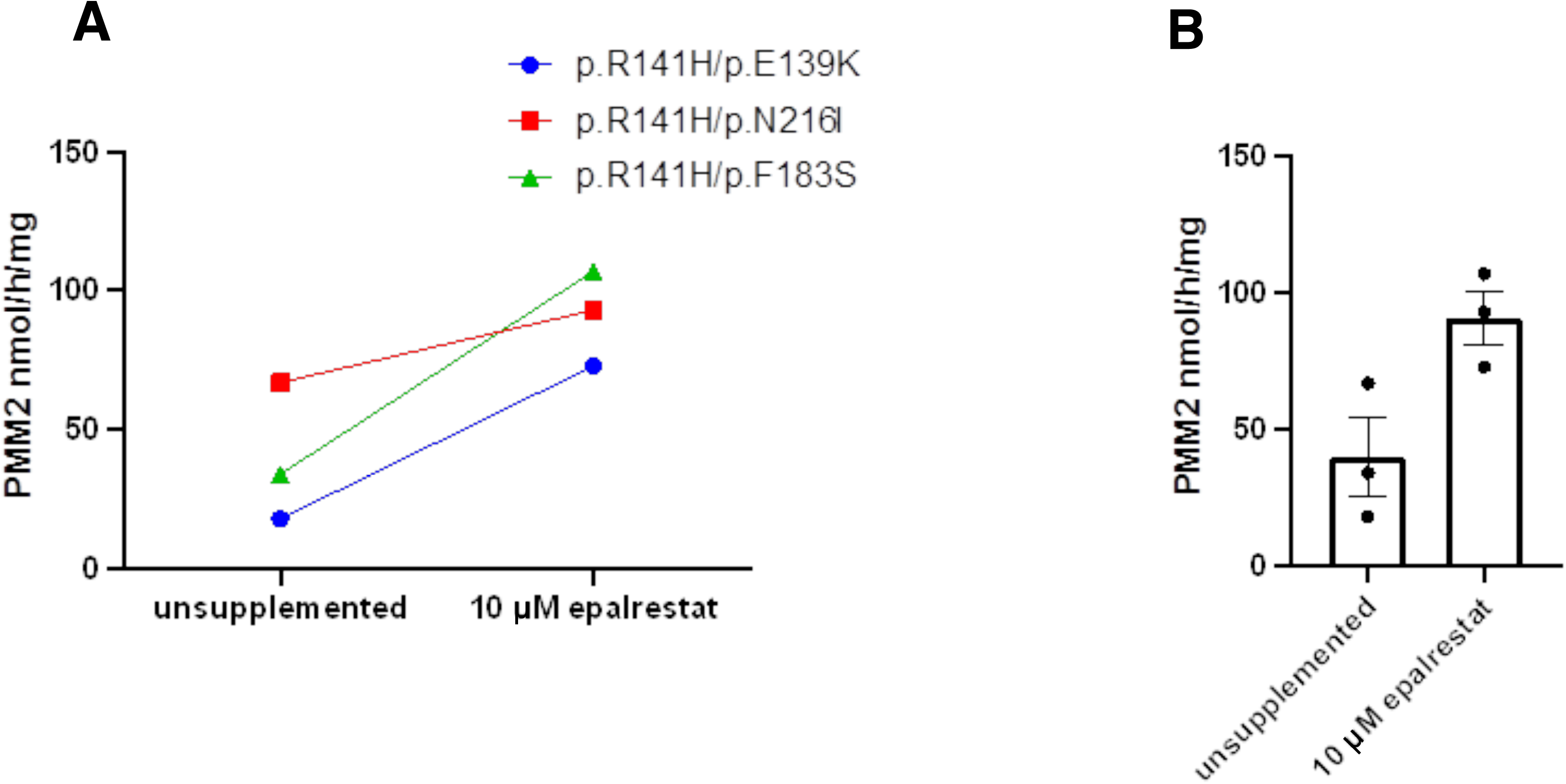
Epalrestat boosts human PMM2 enzymatic activity in multiple PMM2-CDG patient fibroblasts. PMM2 enzymatic activity is expresed as nanomoles per hour per milligram. (**A**) Supplemented samples were treated with 10µM epalrestat for 24 hours. Blue circles are PMM2-CDG fibroblasts with the genotype R141H/E139K Red squares are PMM2-CDG fibroblasts with the genotype R141H/N216I. Green triangles are PMM2-CDG fibroblasts with the genotype R141H/F183S. (**B**) Bar plot of mean PMM2 enzymatic activity with or without epalrestat treatment (p = 0.0321). Each black circle indicates the mean of replicates for each fibroblast. Error bars in the bar graphs indicate standard error of means. P value was determined by t test (T.TEST function in Excel).

**Table 1.** PMM2 enzymatic activity assay of PMM2-CDG patient fibroblast. Supplemented samples were treated with 10µM epalrestat for 24 hours. Technical replicate values from a representative experiment are shown in parentheses).

| PMM2 patient | PMM2 activity (nmol/hr/mg) |  | fold change |
| --- | --- | --- | --- |
|  | unsupplemented | supplemented |  |
| p.R141H/p.E139K | (16; 19; 20)<br>18.33 | (69; 74; 76)<br>73.00 | **4.06** |
| p.R141H/p.N216I | (62; 69; 70)<br>67.00 | (92; 89; 98)<br>93.00 | **1.39** |
| p.R141H/p.F183S | (31; 35; 36)<br>34.00 | (101; 108; 112)<br>107.00 | **3.15** |

Patient R141H/E139K fibroblasts show the lowest un-supplemented PMM2 enzyme activity but attain the largest relative increase in PMM2 enzyme activity levels after 24 hours of epalrestat treatment: 4.06-fold increase from 18 to 72 nmol/h/mg. Patient R141H/N216I fibroblasts have the highest PMM2 enzyme activity baseline and attain the smallest relative increase in PMM2 enzyme activity levels with epalrestat: 1.39-fold increase from 67 to 93 nmol/h/mg. Patient R141H/F183S fibroblasts show robust rescue in epalrestat: 3.15-fold increase from 34 to 107 nmol/h/mg. Averaging across the three genotypes, PMM2 activity increases 2.29-fold over the vehicle-treated baseline (p = 0.0321). Supplemented PMM2 activity levels did not reach the formal patient control reference activity value of >700 nmol/h/mg, yet the identification of several-fold increased PMM2 activity by 10µM epalrestat in 24 hours represents a proof-of-concept for the therapeutic application of epalrestat in remediation of loss of PMM2 enzymatic activity.

### Epalrestat does not increase PMM2 protein abundance

The simplest model to explain how epalrestat increases PMM2 enzyme activity levels is that epalrestat increases PMM2 protein abundance. We demonstrated previously that PMM2 overexpression rescues lethality of yeast models of PMM2-CDG by mass action effects (Lao et al, 2019). The results of immunoblotting with antibody specific to PMM2 clearly show that epalrestat treatment does not increase PMM2 protein abundance in PMM2-CDG fibroblasts or in control fibroblasts (**Figure 7A**).

**Figure 7.**
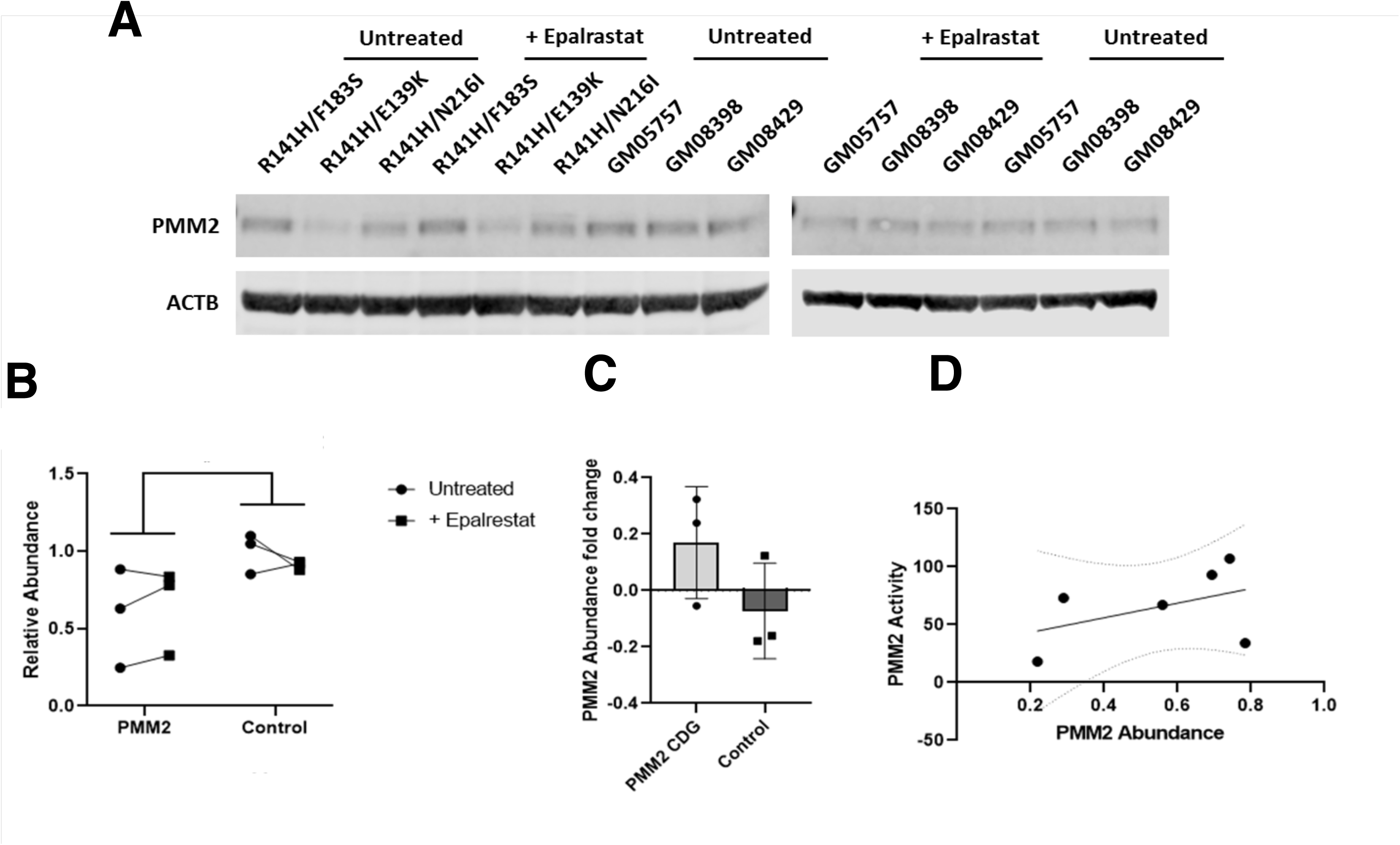
Epalrestat treatment does not increase PMM2 protein abundance. (**A**) PMM2 protein levels determined by immunoblotting in epalrestat-treated PMM2-CDG patient fibroblasts and control fibroblasts. Actin was used a loading control. (**B**) Quantification of PMM2 protein abundance based on band intensity. Black circles represent 10µM epalrestat-treated PMM2-CDG fibroblasts. Black squares represent untreated control fibroblasts. (**C**) PMM2 protein abundance fold change before and after 10µM epalrestat treatment for 24 hours. Black circles represent epalrestat-treated PMM2-CDG fibroblasts. Black squares represent untreated control fibroblasts. (**D**) Plot of PMM2 enzymatic activity as a function of PMM2 protein abundance. Black circles represent three PMM2-CDG patient fibroblast lines and three control fibroblasts.

In two out of three PMM2-CDG fibroblasts, the PMM2 protein abundance increases upon epalrestat treatment, but in the third PMM2-CDG fibroblast PMM2 protein abundance decreases upon epalrestat treatment; and we observed the reverse trend in three control fibroblasts (**Figure 7B**). We averaged PMM2 protein abundance fold change in epalrestat-treated PMM2-CDG fibroblasts and then compared it to the averaged PMM2 protein abundance fold change in control fibroblasts (**Figure 7C**). There is no statistically significant difference. Furthermore, we observed a weak positive trend between baseline PMM2 protein abundance versus baseline PMM2 enzyme activity (**Figure 7D**). Together, these results clearly demonstrate that the increase in PMM2 enzyme activity levels is not explained by increased PMM2 protein abundance. The increase in PMM2 enzyme activity levels must therefore be explained by increased catalysis of each PMM2 homodimer already present in the cell, not an increase in the total cellular concentration of PMM2 protein via transcriptional upregulation of PMM2 mRNA.

### Epalrestat increases PMM2 mRNA expression and rescues ER stress markers in worms

It was recently reported that a *pmm2* hypomorphic mutant zebrafish constitutively activates NRF2 and furthermore that the small molecule NRF2 activator sulforaphane ameliorates endoplasmic reticulum (ER)-associated stress markers (Mukaigasa et al, 2018). Therefore, we tested whether epalrestat acts similarly to sulforaphane by interrogating mRNA expression levels of the worm orthologs of the ER stress markers that are induced in a *pmm2* hypomorphic mutant zebrafish. The results of this qPCR analysis are shown in **Supplemental Figure 5**. *PMM-2* transcript levels are increased by 15µM epalrestat. Transcript levels of *HSP-1*, which encodes a heat shock protein, are also increased by epalrestat treatment. Conversely, the ER stress markers *IRE-1*, *PEK-1*, *SKN-1*, and *GCS-1*, which are constitutively elevated in the F119L mutant, are modestly decreased by treatment with epalrestat. These results bolster confidence in the translational fidelity of our F119L worm model of PMM2-CDG because it phenocopies the constitutively active ER stress response seen in a zebrafish model of PMM2-CDG.

## Discussion

We set out in this study to build on the results of our previous yeast PMM2-CDG disease modeling and drug repurposing efforts (Lao et al, 2019). We filled a gap in PMM2-CDG disease models by generating a nematode model based on a specific PMM2-CDG variant, in this case the second-most common missense allele F119L (F125L in worms). We confirmed that the F119L nematode model has between 30-40% residual *PMM-2* enzymatic activity, comparable to a haploid F119L yeast model. We identified hypersensitivity to the proteasome inhibitor bortezomib as a growth-based phenotype in an unbiased whole-organism drug repurposing screen. A comparative analysis of yeast and worm PMM2 repurposing hits revealed overlap in one structural class: plant-based polyphenols. Using a PMM2 enzymatic activity assay in PMM2-CDG patient fibroblasts, we showed that several yeast and worm repurposing candidates increase PMM2 enzymatic activity. These compounds are the first known small molecule activators of PMM2 enzymatic activity.

Analysis of structure-activity relationships and cross-species testing suggested that plant-based polyphenols are acting as aldose reductase inhibitors (ARIs). Upon testing commercially available ARIs, we showed conclusively that epalrestat boosts PMM2 enzymatic activity in both nematodes and PMM2-CDG patient fibroblasts. Epalrestat is the only ARI approved for the treatment of diabetic neuropathy in humans and has been used safely for nearly three decades. It has been used to treat peripheral neuropathy in geriatric diabetic patients in Asia for several decades. We believe our results justify repurposing epalrestat for PMM2-CDG, starting with a small safety and efficacy study involving up to ten PMM2-CDG patients. The only other example of drug repurposing for PMM2-CDG involves the carbonic anhydrase inhibitor acetazolamide, and the therapeutic thesis is that acetazolamide will treat the cerebellar syndrome of PMM2-CDG but not correct the root cause of disease which is PMM2 enzyme insufficiency (Martinez-Monseny et al, 2019).

What is the mechanism of action by which epalrestat potentiates PMM2 enzymatic activity? We initially hypothesized that activation of NRF2 would lead to increased PMM2 mRNA levels, which in turn would lead to increased PMM2 protein abundance. Consistent with that hypothesis, treatment of *pmm2* zebrafish mutant with the NRF2 activator sulforaphane resulted in rescue of phenotypic defects (Mukaigasa et al, 2018). We know from our studies of yeast PMM2-CDG models that overexpression of F119L mutant PMM2 protein can rescue growth by mass action effects (Lao et al, 2019). Four plant-based polyphenols – fisetin, rhamnetin, pyrogallin and purpurogallin-4-carboxylic acid – are active in the Keap1-Nrf2 cellular reporter assay, but only two of them – fisetin and pyrogallin – boost PMM2 enzymatic activity. Several reports indicate that epalrestat activates NRF2 in cultured cells (Yama et al, 2016; Yama et al, 2015). Using quantitative RT-PCR, we showed that epalrestat treatment of *pmm-2* F125L/F125L homozygote worms modestly upregulates *SKN-1* transcripts. *SKN-1* is the worm ortholog of NRF2. As a direct test of this model, we quantified PMM2 protein abundance in PMM2-CDG patient fibroblasts by immunoblotting. Epalrestat does not increase PMM2 protein abundance in any of the fibroblast lines tested.

Therefore, we favor a model where epalrestat acts post-translationally to boost PMM2 enzymatic activity by preventing glucose from being shunted down the polyol pathway. Aldose reductase is the first biosynthetic enzyme in the polyol pathway, and ARIs block the conversion of glucose to sorbitol. Excess glucose would then favor the production of glucose-1,6-bisphosphate, an endogenous small molecule coactivator of PMM2 (Pirard et al, 1999; Monticelli et al, 2019). At the same time, inhibition of the polyol pathway reduces the levels of advanced glycation end products, which would attenuate the sequelae of ER stress and oxidative stress. In humans, epalrestat is known to reduce the levels of carboxymethyl lysine, a known advanced glycation end product (Kawai et al, 2010). It is also possible that there is a complex interaction between activation of NRF2 and inhibition of the polyol pathway, the former acting at the transcriptional level and the latter acting post-translationally.

Our model make several testable predictions. First, the concentration of glucose-1,6-bisphosphate will increase in yeast, worms and human cells treated with epalrestat. Second, the concentration of sorbitol will decrease in yeast, worms and human cells treated with epalrestat. Third, CRISPR knockout of aldose reductase in PMM2-CDG patient fibroblasts will block epalrestat’s activation of PMM2 enzyme activity. In the interim, decades of safe and effective administration of epalrestat in geriatric populations for peripheral neuropathy suggests that the drug may also be safe in pediatric settings necessitated by PMM2-CDG. Peripheral neuropathy is observed in almost all PMM2-CDG patients, so a therapeutic rationale for repurposing epalrestat for PMM2-CDG is compelling.

There are over 180 publications on the use of epalrestat in the past 25 years in the peer-reviewed literature. Epalrestat has been studied in three pivotal registration studies, and multiple post market studies have been conducted. The drug has an excellent safety profile, with the most common side effects reported being nausea, vomiting, diarrhea, and elevated liver enzymes, and the most severe side effect reported being liver failure.

Will the PMM2 enzyme activity level increases in epalrestat-treated patient fibroblasts lead to clinical benefit in patients? The expectation is that epalrestat will treat the peripheral neuropathy of PMM2-CDG patients but will it have impact beyond peripheral neuropathy? Based on Mayo Clinic’s PMM2-CDG biobank, 55 affected fibroblast lines exhibit a range of PMM2 enzymatic activity between 18-307 nmol/hr/mg. Mayo Clinic uses a diagnostic cutoff at 307 nmol/hr/mg. 66 unrelated non-carrier control fibroblast lines exhibit a range of PMM2 enzymatic activity between 712-2066 nmol/hr/mg. Mayo Clinic reports as healthy PMM2 enzymatic activity as any value > 712 nmol/hr/mg. It is believed by clinicians working on PMM2-CDG that > 100 nmol/hr/mg of PMM2 enzymatic activity would translate to clinical benefit.

## Acknowledgements

We acknowledge Maggie’s PMM2-CDG Cure as a funding source, and we thank the anonymous reviewers for feedback that significantly improved the manuscript.

---

**Supplementary Figure 1.**
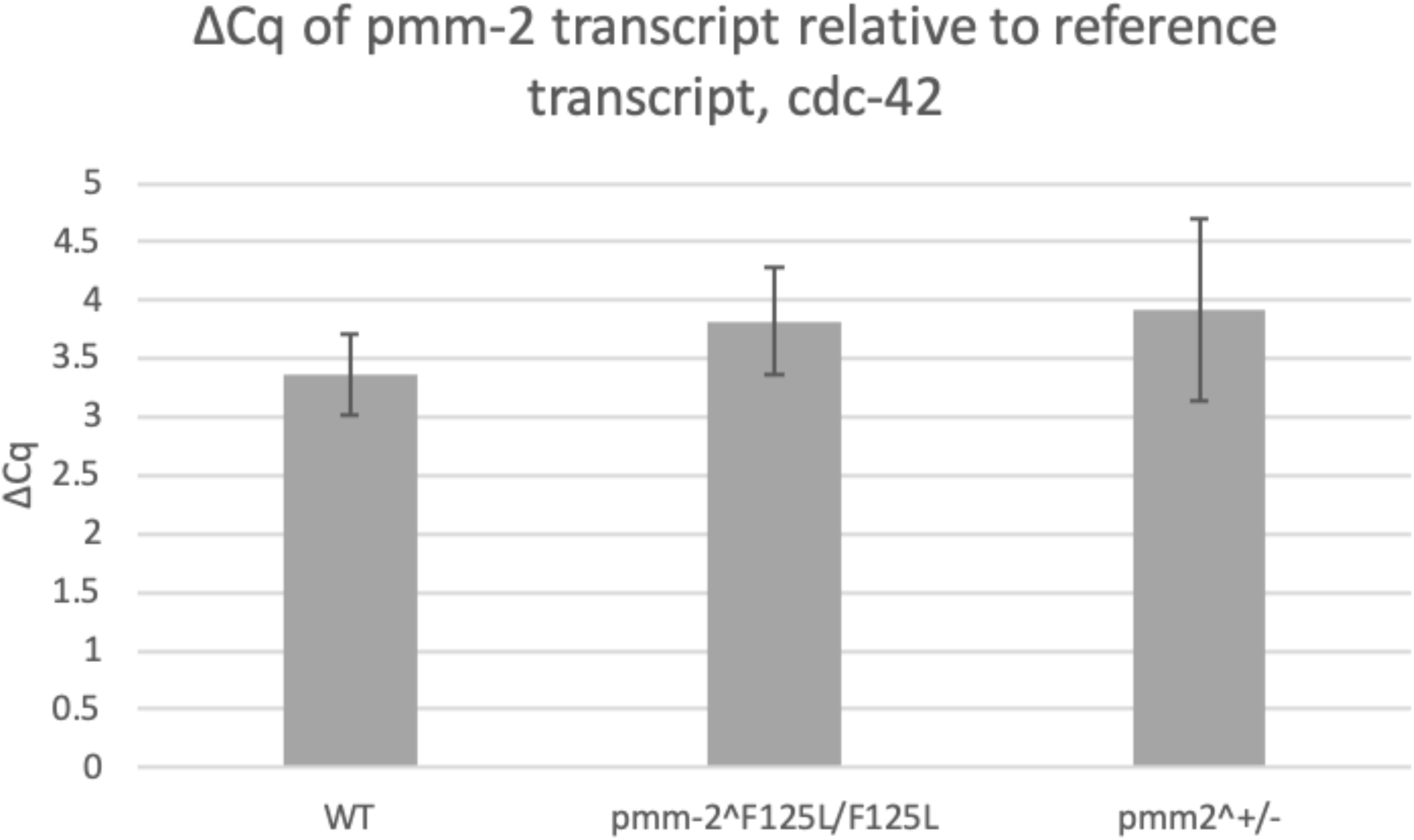
Quantitative RT-PCR analysis of PMM-2 expression in pmm-2 F125L (F119L) homozygote mutant worms compared to the heterozygous *pmm-2* deletion mutant and a wildtype reference strain (N2). ΔCq values were calculated by subtracting the reference transcript (cdc-42) cycle quantification (Cq) number from that of the target transcript (pmm-2). From left to right, ΔCq values for N2, *pmm-2* F125L/F125L homozygote mutant (COP1626) and pmm-2 heterozygous mutant (VC3054) are displayed. Each bar consists of data from three biological replicates. Error bars are standard error of mean ΔCq across three replicates ΔCq values are not significantly different from wildtype indicating that both homozygous and heterozygous mutants produce the same level of *pmm-2* transcript as N2.

**Supplementary Figure 2.**
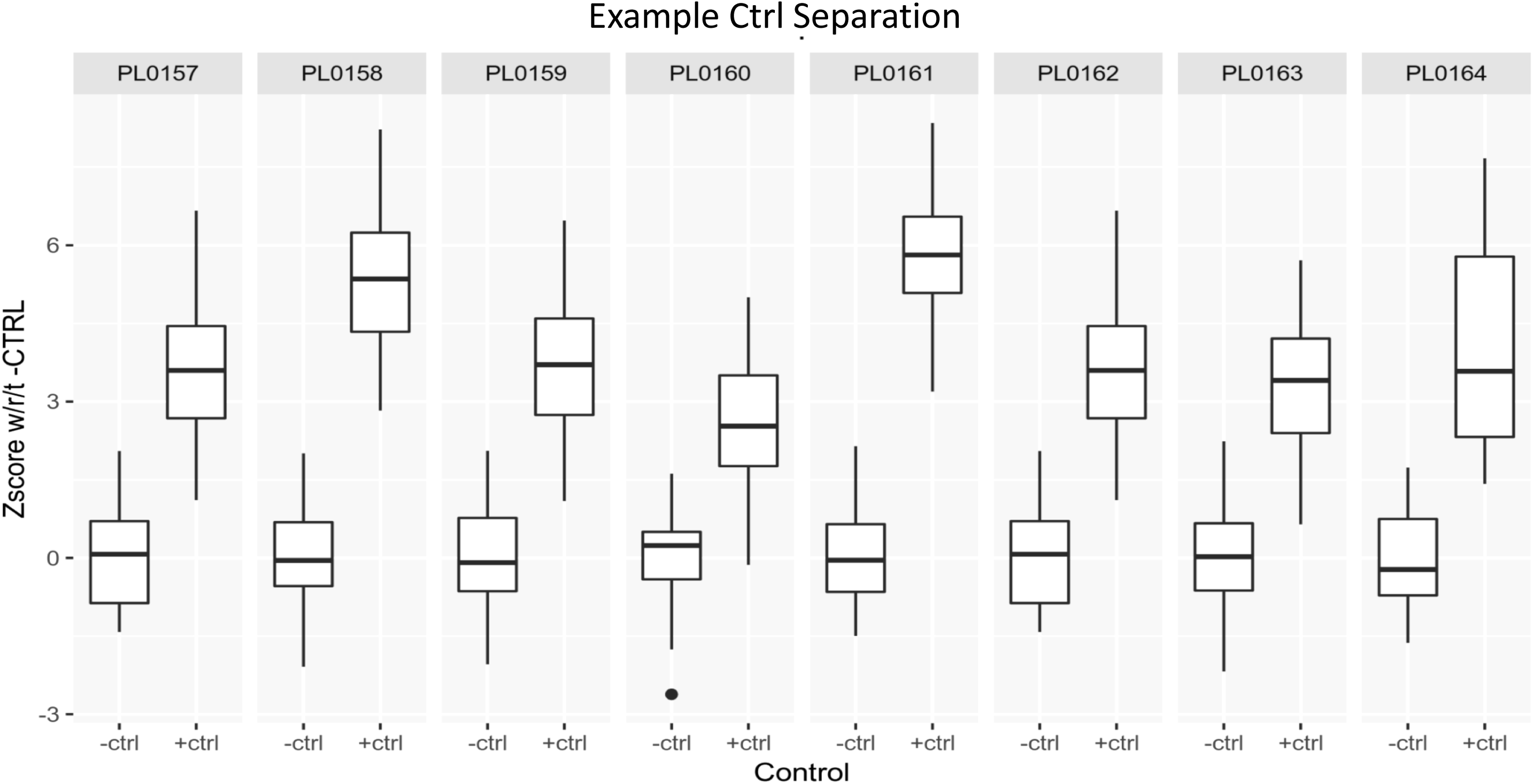
Box and whisker plots of z scores of positive and negative control wells from a representative replicate of the Microsource Spectrum drug repurposing library screen. Negative controls have Z score values of 0, whereas Z score values of positive controls are > 2.

**Supplementary Figure 3.**
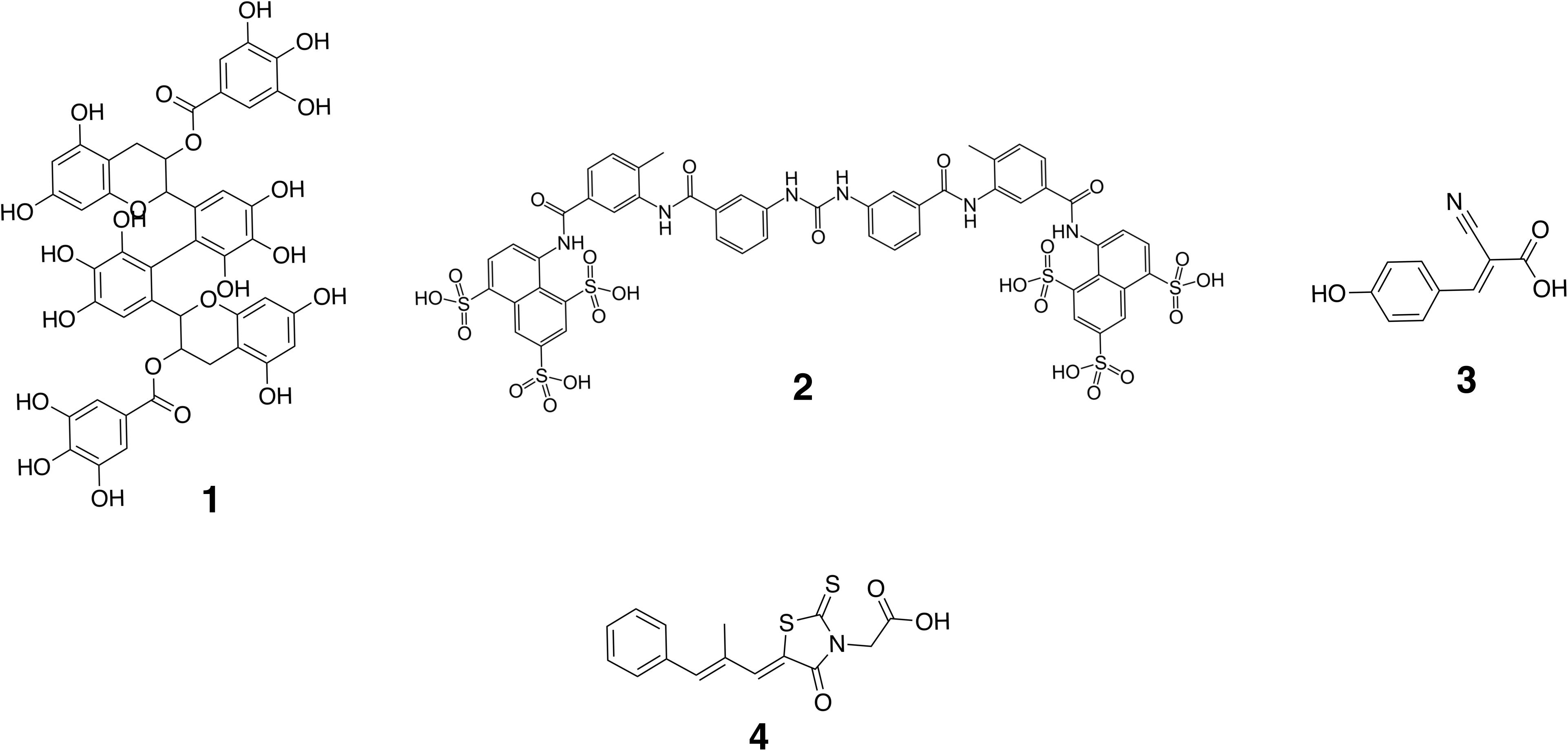
Chemical structures of yeast repurposing candidates from Lao et al, 2019 (**1**) 2’-2’-bisepigallocatechin digallate (**2**) suramin (**3**) alpha-cyano-4-hydroxycinnamic acid, and the chemical structure of the aldose reductase inhibitor epalrestat (**4**). Notice the shared carboxylic moiety of **3** and **4**.

**Supplementary Figure 4.**
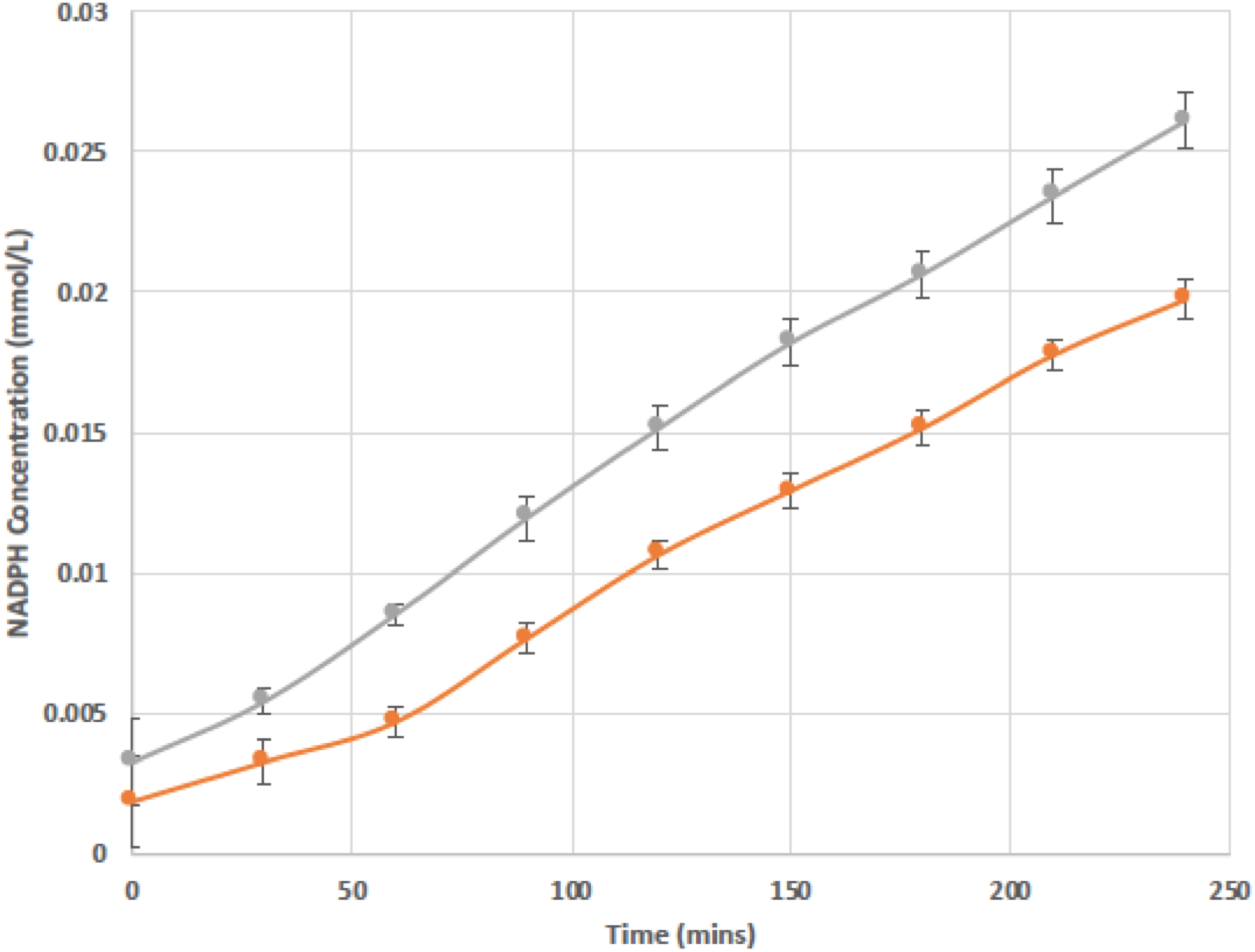
PMM2 enzymatic activity assay of R141/F119L PMM2-CDG patient fibroblasts. Supplemented samples were treated with 10µM epalrestat for 24 hour. Error bars in the bar graphs indicate standard error.

**Supplementary Figure 5.**
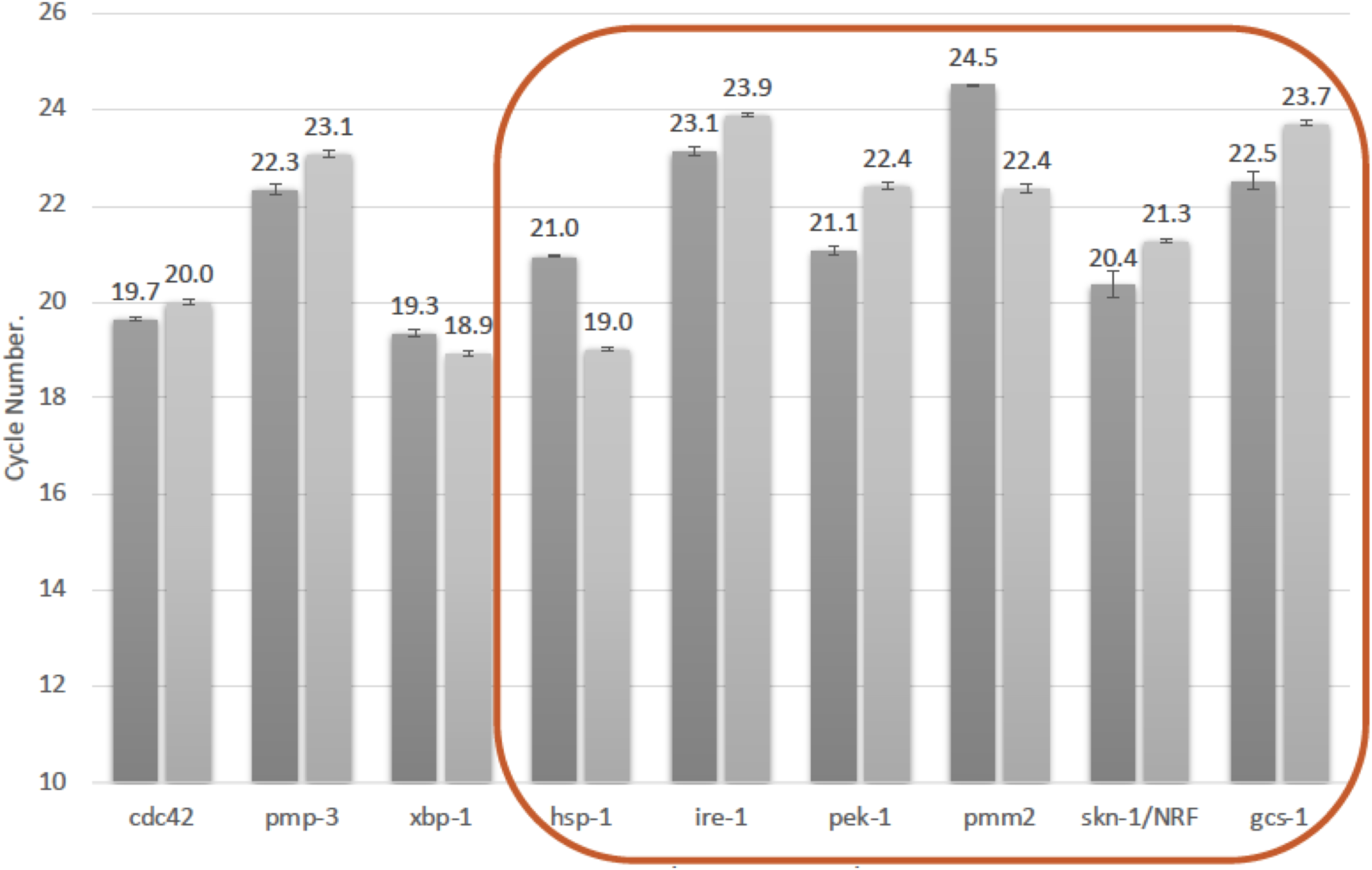
Quantitative RT-PCR analysis of PMM-2 and ER stress marker expression in *pmm-2* F125L/F125L homozygote mutant worms.

**Supplementary Figure 6.**
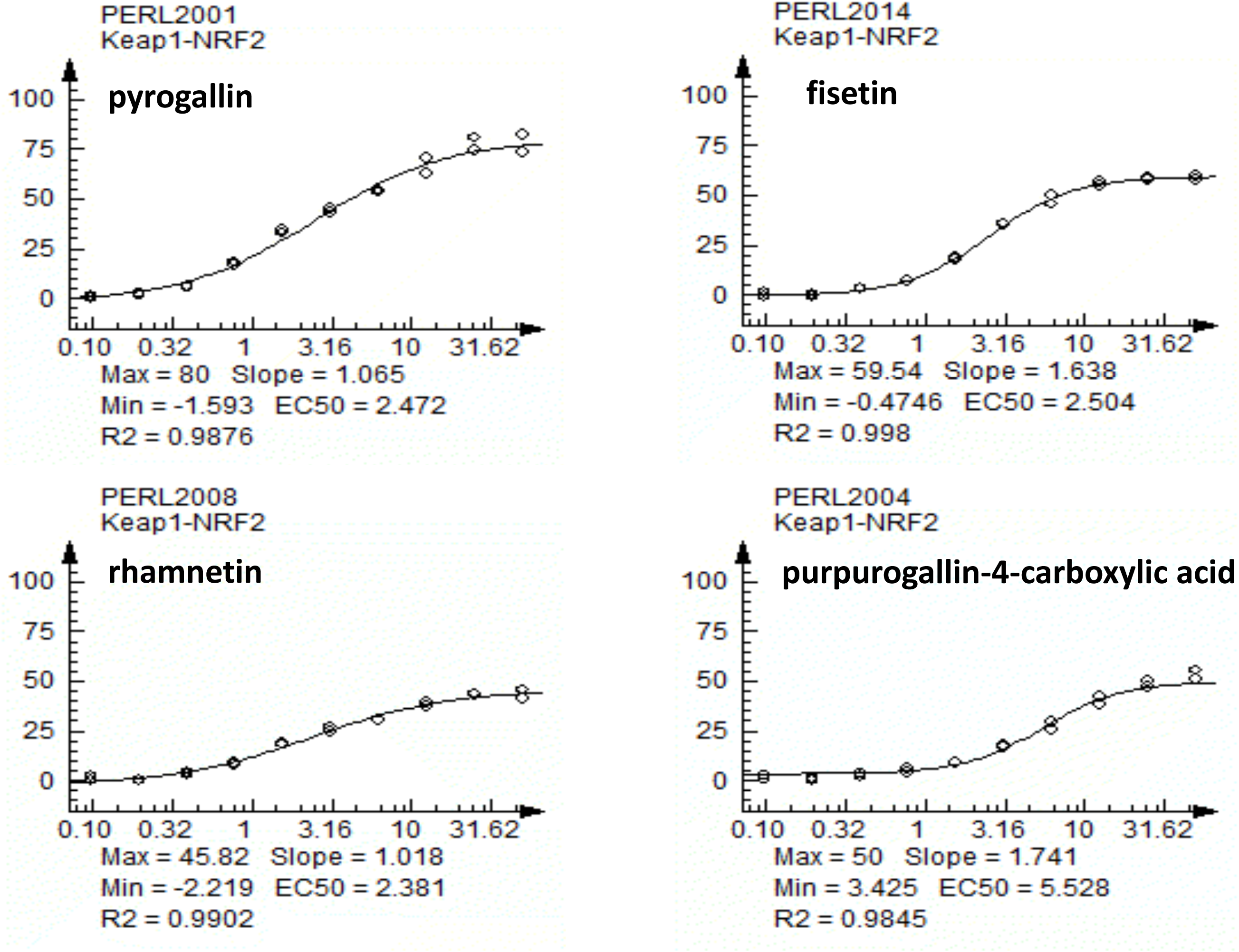
Dose-response curves of four worm PMM2 repurposing hits in a Keap1-NRF2 reporter activation assay in human cells generated by DiscoverX. (**A**) pyrogallin. (**B**) fisetin. (**C**) rhamnetin. (**D**) purpurogallin-4-carboxylic acid.

**Supplementary Figure 7.**
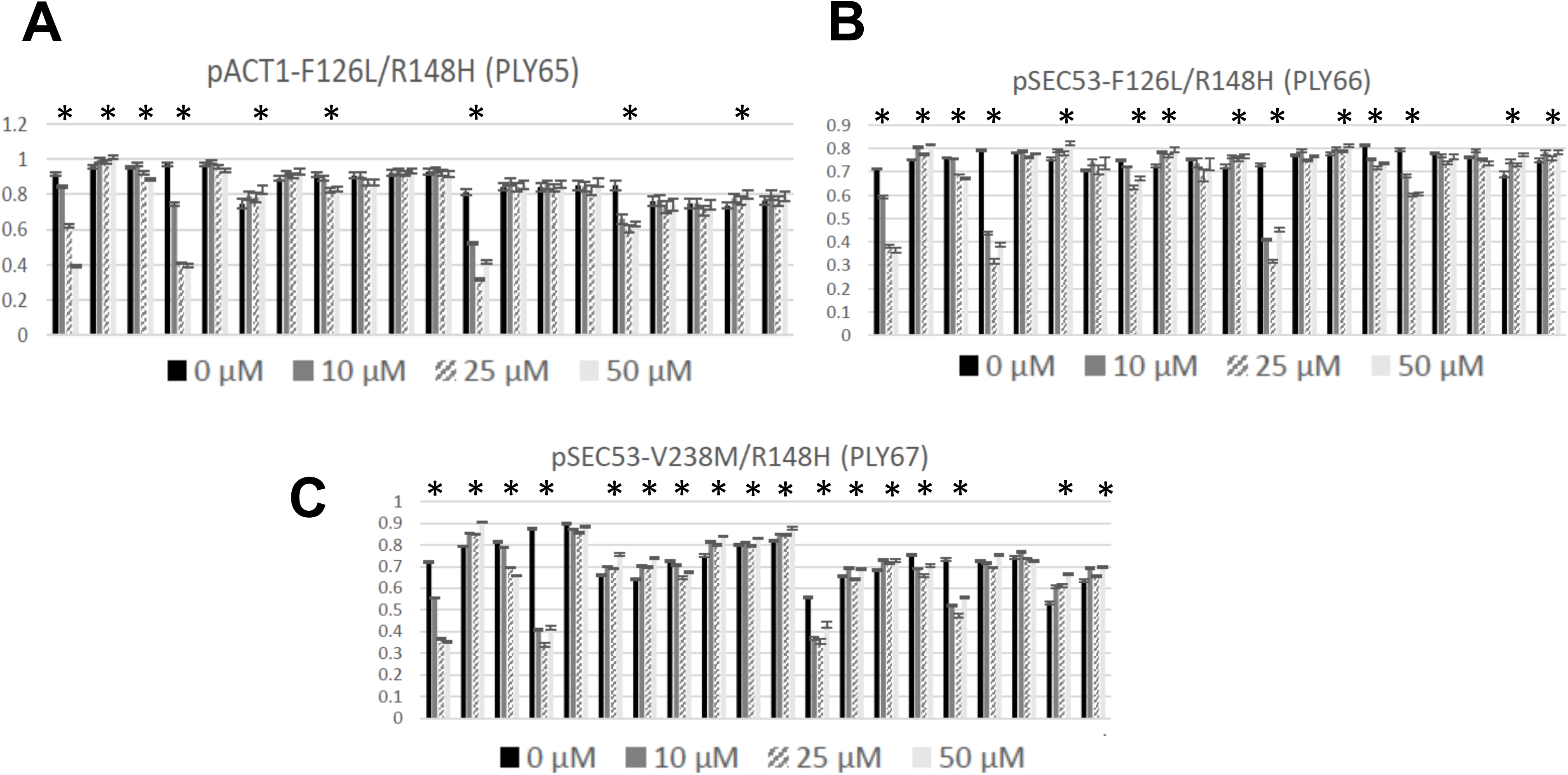
Growth of three yeast PMM2-CDG models (pACT1-F126L/R148H; pSEC53-V238M/R148H; pSEC53-F126L/R148H) in the presence of the 20 worm repurposing candidates at 10µM, 25µM and 50µM. Compounds tested from left to right: pyrogallin, amidol, baicalein, purpurogallin-4-carboxylic acid, gossypetin, quercetin tetramethylether, 3-methoxycatechol, rhamnetin, theaflavin monogallate, hieracin (tricetin), epicatechin monogallate, 3,4-didesmethyl-5-deshydroxy-3-ethoxyschleroin, 2,3,4-trihydroxy-4-methoxybenzophenone, koparin, fisetin, edaravone, ellagic acid, levodopa, dobutamine and ethylnorepinephrine. Asterisks indicate compounds that have either positive or negative effects on growth.

**Supplementary Figure 8.**
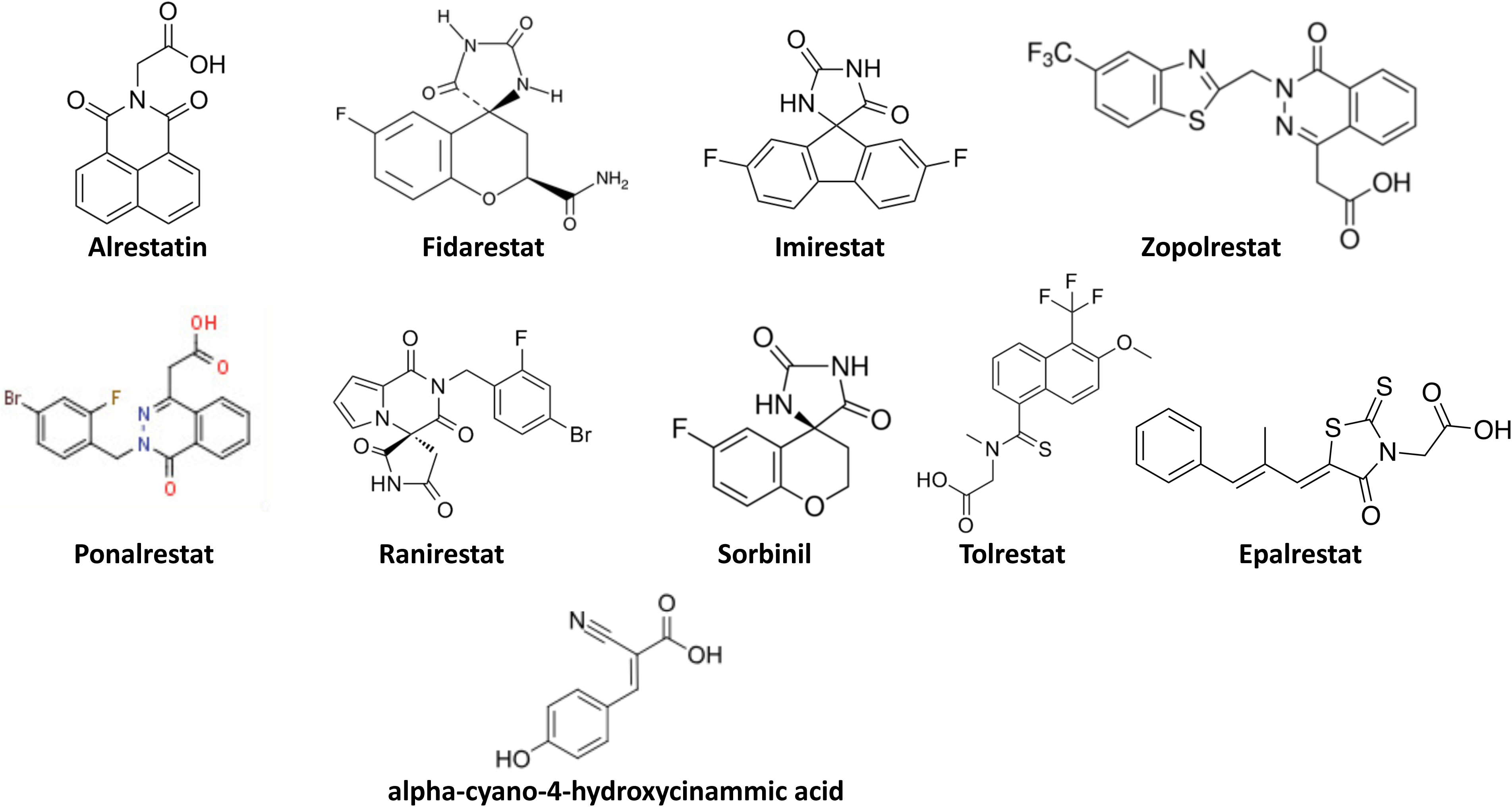
Chemical structures of 10 commercially available aldose reductase inhibitors tested in PMM2-CDG R141H/F119L fibroblasts.

## Notes

#### Summary of Updates

This revised manuscript (v2.0) was significantly revised and improved during peer review at Disease Models & Mechanisms.

